# Chloroplast ABC peptide transporters TAP1, NAP8, and ATH12 are essential for heat-induced peptide export and play a key role in thermotolerance in *Arabidopsis thaliana*

**DOI:** 10.64898/2026.03.05.709749

**Authors:** Nicolaj Jeran, Guido Domingo, Luca Tadini, Elena Costantini, Cecilia Lasorella, Stefania Fortunato, Milena Runge, Chiara Bertaso, Federico Vita, Maria Concetta De Pinto, Candida Vannini, Paolo Pesaresi

## Abstract

ATP-binding cassette (ABC) transporters mediate substrate translocation across membranes by using energy from ATP hydrolysis. While ABC peptide exporters have been characterized in the mitochondria of metazoans and yeast, corresponding chloroplast peptide transport systems in plants remain uncharacterized. Using *in silico* and experimental approaches, we identify three previously uncharacterized *Arabidopsis thaliana* ABCB half-transporters - TAP1, NAP8, and ATH12 - that localize to the chloroplast inner envelope and form homodimers. These proteins are phylogenetically related to known peptide exporters, and functional complementation in *Saccharomyces cerevisiae* demonstrates that each plant transporter can rescue the heat sensitivity of a *Δmdl1* mutant, indicating conserved peptide export activity. In chloroplast peptide efflux assays, peptide export upon heat stress was strongly reduced only in the *tap1 nap8 ath12* triple mutant, but not in single or double mutants, indicating functional redundancy. Furthermore, mass spectrometry of chloroplast supernatants revealed an abundance of thylakoid-derived hydrophilic peptides in the wild type, predicted to have antioxidant activity. Under heat stress, the triple mutant displayed increased sensitivity, characterized by reduced biomass, chlorophyll and carotenoid content, and compromised photosynthetic efficiency. Comprehensive analyses revealed altered redox homeostasis in the triple mutant, including modified antioxidant dynamics, differential antioxidant enzyme activities, and distinct gene expression profiles compared with the wild type. Our findings demonstrate that TAP1, NAP8, and ATH12 constitute a chloroplast peptide export system required for efficient peptide release under heat stress, with a role in *Arabidopsis* thermotolerance. These results provide new insights into organellar peptide transport and its integration with stress mitigation mechanisms in plants.

## Introduction

Peptide transporters are ubiquitous membrane proteins that mediate the ATP-dependent translocation of oligopeptides across biological membranes and play essential roles in nutrient acquisition, protein turnover, and cellular stress responses (Rees et al., 2009; Thomas and Tampé, 2020). Among these systems, ATP-binding cassette (ABC) peptide transporters constitute one of the most ancient and evolutionary conserved transporter families, present across all domains of life. Their widespread distribution underscores the central importance of peptides not only as metabolic substrates but also as signalling molecules (Davidson et al., 2008; Locher, 2016). In bacteria, peptide transport is predominantly mediated by ABC importers, including the well-characterized Dpp, Tpp, and Opp (<u>D</u>ip<u>p</u>eptide, <u>T</u>ri<u>p</u>eptide, <u>O</u>ligo<u>p</u>eptide <u>p</u>ermeases) systems, which enable the uptake of di-, tri-, and oligopeptides to sustain growth, nitrogen metabolism, and quorum sensing (Detmers et al., 2001; Monnet, 2003). Beyond nutrient acquisition, bacterial peptide transporters contribute to virulence, antibiotic resistance, and environmental sensing, highlighting their multifunctional roles in microbial physiology (Lewis et al., 2012; Abele and Tampé, 2018). In metazoans, ABC peptide transporters are primarily involved in intracellular peptide trafficking, most notably through the transporter associated with antigen processing (TAP1/TAP2), which mediates the ATP-dependent translocation of proteasome-derived peptides into the endoplasmic reticulum for MHC class I antigen presentation (Nijenhuis and Hämmerling, 1996; Lehnert et al., 2016). Collectively, these studies establish peptide transporters as integral components of cellular homeostasis and signalling networks, extending their function far beyond simple metabolite exchange.

Accumulating evidence further highlights a pivotal role for peptide transporters in intracellular communication, particularly in organelle-derived stress signalling pathways (Quirós et al., 2016). Mitochondria continuously degrade proteins through ATP-dependent proteases, generating peptides that must be efficiently exported both to prevent their toxic accumulation within the organelle and to convey information about mitochondrial functional status to the rest of the cell (Lebeau et al., 2018). In *Caenorhabditis elegans*, the inner mitochondrial membrane ABC transporter HAF-1 mediates the export of peptides generated by mitochondrial proteolysis into the cytosol and is essential for activation of the mitochondrial unfolded protein response (UPR^mt^; Haynes et al., 2010). A functionally analogous system operates in yeast, where the *Saccharomyces cerevisiae* ABC transporter Mdl1 exports peptides produced by the m-AAA protease, thereby sustaining mitochondrial proteostasis and respiratory competence (Young et al., 2001; Arnold et al., 2006). Additional roles for Mdl1 have been reported, including contributions to resistance against oxidative stress (Chloupková et al., 2003) and to the cellular response to clozapine, an antipsychotic drug (Theron et al., 2024). Despite being identified in distinct model organisms, *Ce*HAF-1 and *Sc*Mdl1 exemplify a conserved mechanism in which peptide export from the inner mitochondrial membrane, mediated by homodimeric ABC transporters, links organellar proteolysis to nuclear gene expression via retrograde signalling pathways (Fiorese et al., 2016). Consistent with this model, mitochondria isolated from *haf-1* and *Δmdl1* mutants released fewer peptides into the incubation buffer than wild-type mitochondria in the presence of ATP. Notably, the corresponding mutant strains exhibited increased sensitivity to elevated temperatures, a major stress that challenges cellular proteostasis (Young et al., 2001; Haynes et al., 2010; Jarolim et al., 2013).

In plants, by contrast, the contribution of peptide transporters to organelle-to-nucleus retrograde signalling under heat stress remains poorly understood, despite the presence of numerous ABC transporters localized to mitochondria and chloroplasts (Kang et al., 2011; Hwang et al., 2016). Heat stress responses in plants are tightly linked to the maintenance of redox homeostasis (Hendrix et al., 2023) and impose severe challenges to protein folding and stability, frequently resulting in protein misfolding and aggregation (Staacke et al., 2025). To mitigate proteotoxic damage, plants induce a suite of molecular chaperones that are specifically upregulated under heat and other stress conditions. Among these, the chloroplast-localized unfoldase CLPB3 plays a central role in resolving stromal protein aggregates and is strongly induced at elevated temperatures (Myouga et al., 2006; Parcerisa et al., 2020; Kreis et al., 2023; Jeran et al., 2025). In parallel, chloroplasts harbour a highly elaborate and compartmentalized proteolytic network that ensures protein quality control and regulated protein turnover, particularly under adverse environmental conditions such as heat stress (van Wijk, 2015; Nishimura et al., 2017; van Wijk, 2024). The stromal CLP protease complex constitutes the major ATP-dependent protease responsible for bulk protein degradation, whereas thylakoid-associated FTSH and DEG proteases are key players in photosystem II repair and selective protein turnover. Additional proteolytic activities have been described at the chloroplast inner envelope membrane; however, these systems remain comparatively less well characterized. By analogy with mitochondrial systems, it is plausible that peptides generated during chloroplast proteolysis under heat stress are actively exported to preserve organelle functionality (Nishimura et al., 2017), raising the possibility that specific ABC transporters function as peptide exporters linking chloroplast proteostasis to nuclear gene expression. Support for a proteolysis-dependent retrograde signalling mechanism is provided by studies on singlet oxygen (¹O₂)-mediated signalling, particularly the EXECUTER1 (EX1) pathway (Wagner et al., 2004; Liu et al., 2024). EX1 is rapidly degraded by the FTSH2 protease following ¹O₂ production at photosystem II, and this proteolytic event is essential for downstream nuclear transcriptional reprogramming and stress responses (Dogra et al., 2019). The requirement for EX1 degradation supports the concept that proteolysis-derived peptides act as mobile signalling intermediates in chloroplast retrograde pathways (Zhao et al., 2025).

Here, we identify three chloroplast-localized ABC peptide transporters - TAP1, NAP8, and ATH12 - that mediate ATP-dependent peptide export from chloroplasts during heat stress. We show that their combined loss impairs peptide efflux, redox homeostasis, and heat tolerance in Arabidopsis.

## Results

### *In silico* identification of plastid-localized ABC peptide transporters in *Arabidopsis thaliana*: TAP1, NAP8, and ATH12

To identify putative plastid-localized peptide transporters *in silico*, the amino acid sequences of the mitochondrial peptide exporters *Ce*HAF-1 and *Sc*Mdl1 were used as queries to search the *Arabidopsis thaliana* reference protein database. These searches retrieved 87 and 80 significant protein hits (E-value < 0.05) for the *Ce*HAF-1 and *Sc*Mdl1 queries, respectively (Table S1), all of which were annotated as members of the ABC transporter family. To refine the candidate list, significant hits were further filtered by applying a BLAST score threshold greater than 100 and by assessing predicted subcellular localization. This two-step filtering procedure reduced the dataset to three previously uncharacterized proteins that satisfied both criteria: Transporter Associated with Antigen Processing 1 (TAP1; also known as ABCB26; encoded by *AT1G70610*), Non-Intrinsic ABC Protein 8 (NAP8; also known as ABCB28 or TAP-related protein 1; encoded by *AT4G25450*), and ABC Two Homolog 12 (ATH12; also known as ABCB29; encoded by *AT5G03910*).

The ABC protein superfamily is subdivided into eight subfamilies based on domain organization, membrane topology, and phylogenetic relationships (Sánchez-Fernández et al., 2001; Verrier et al., 2008; Kang et al., 2011). To classify TAP1, NAP8, and ATH12 within this framework, their amino acid sequences were analysed for domain composition and the presence of transit peptides. For comparison, the sequences of *Ce*HAF-1 and *Sc*Mdl1 were included in all analyses. Whereas *Ce*HAF-1 and *Sc*Mdl1 possess a predicted mitochondrial transit peptide (mTP; Fig. 1A, in red), all three Arabidopsis candidates were predicted to harbour an N-terminal chloroplast transit peptide (cTP; Fig. 1A, in green). In addition, each protein exhibited a conserved half-transporter architecture, comprising a single N-terminal transmembrane domain (TMD; Fig. 1A, in blue) with six predicted α-helices (yellow boxes) followed by a C-terminal nucleotide-binding domain (NBD; Fig. 1A, in orange). These features support the assigment of TAP1, NAP8 and ATH12 to the ABCB half-transporter subfamily, in agreement with previous genome-wide classifications of Arabidopsis ABC proteins(Sánchez-Fernández et al., 2001; Kang et al., 2011).

**Figure 1.**
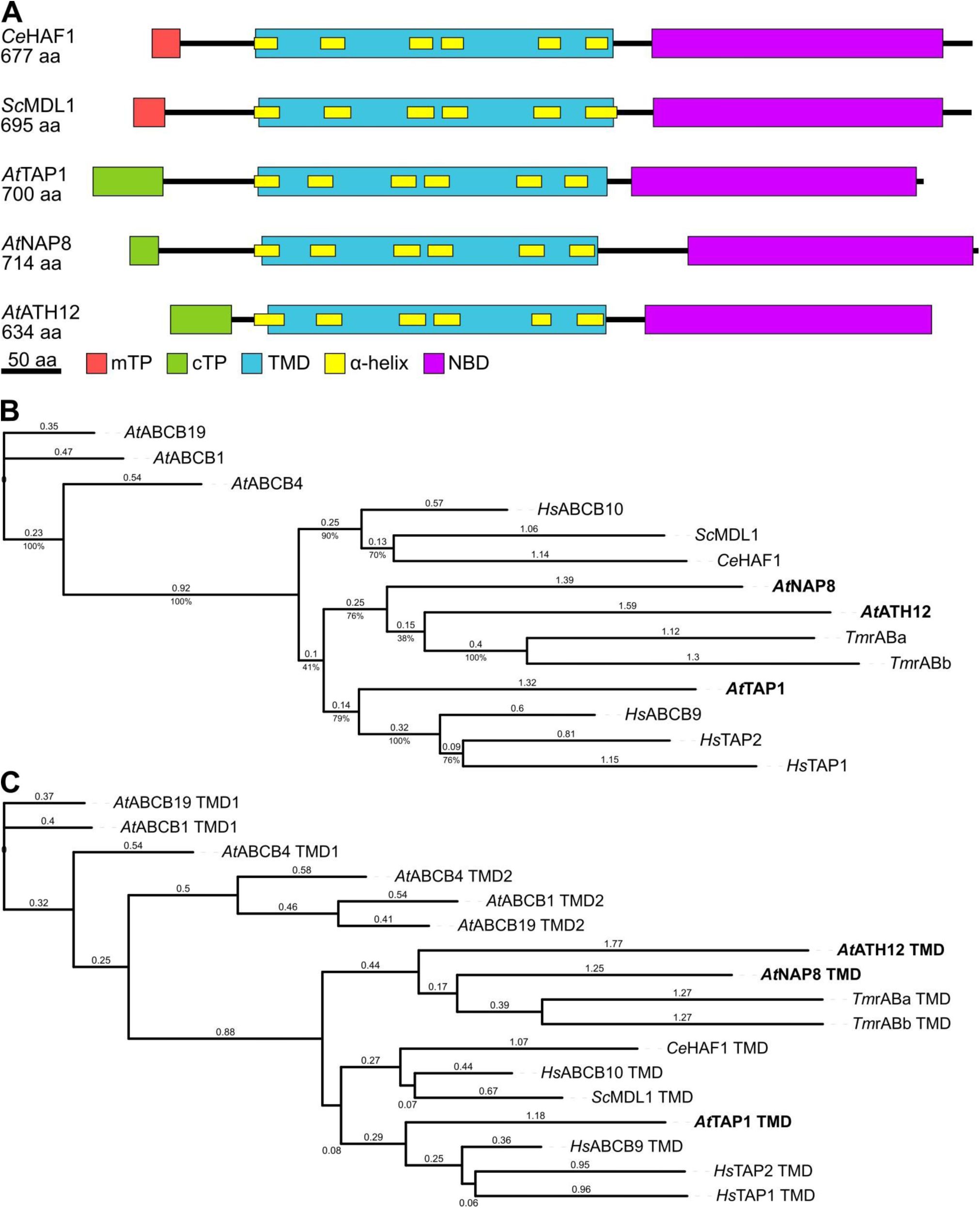
TAP1, NAP8, and ATH12 phylogenetically associate with characterized ABC peptide transporters. **A)** Schematic representation of the domain architecture of the indicated ABC transporters. Predicted chloroplast transit peptides (cTP, green), mitochondrial transit peptides (mTP, red), transmembrane domains (TMD, blue), nucleotide-binding domains (NBD, purple), and individual transmembrane helices (yellow) are indicated. All protein models are drawn to scale and aligned at the first amino acid of the first predicted transmembrane helix. **B)** Phylogenetic tree based on multiple sequence alignment (MSA) of full-length protein sequences from the indicated transporters. Arabidopsis ABCB auxin transporters *At*ABCB1, *At*ABCB4, and *At*ABCB19 were included as an outgroup. Candidate Arabidopsis plastid peptide transporters are highlighted in bold. Bootstrap support values (percent) were calculated from 100 replicates; branch lengths represent phylogenetic distances. Species abbreviations: *Tm*, *Thermus thermophilus*; *Sc*, *Saccharomyces cerevisiae*; *At*, *Arabidopsis thaliana*; *Ce*, *Caenorhabditis elegans*; *Hs*, *Homo sapiens*. **C)** Phylogenetic tree derived from an MSA of the transmembrane domains (TMDs) only, demonstrating that the observed clustering is independent of nucleotide-binding domain conservation.

To further substantiate the homology between the Arabidopsis candidates and functionally characterized peptide exporters, phylogenetic analyses were performed using multiple sequence alignments of TAP1, NAP8, and ATH12 together with eight established peptide transporters from bacteria, yeast, and metazoans (Nijenhuis and Hämmerling, 1996; Young et al., 2001; Wolters et al., 2005; Zhao et al., 2008; Haynes et al., 2010; Liesa et al., 2012; Lehnert et al., 2016; Nöll et al., 2017). Three Arabidopsis ABCB auxin transporters were included as an outgroup to provide phylogenetic context (Noh et al., 2001; Lin and Wang, 2005; Santelia et al., 2005). In the resulting phylogenetic tree (Fig. 1B), the auxin transporters formed a clearly distinct clade, indicating limited sequence similarity to peptide transporters. In contrast, all characterized peptide transporters clustered together with TAP1, NAP8, and ATH12 within a single major branch, supporting their evolutionary relatedness. Within this branch, TAP1 grouped with the metazoan peptide transportersTAP1/ABCB2, TAP2/ABCB3, and TAPL/ABCB9, whereas NAP8 and ATH12 clustered with the bacterial peptide exporters *Tmr*AB from *Thermus thermophilus*. The mitochondrial peptide exporters *Sc*Mdl1, *Ce*HAF-1, and ABCB10 from yeast, nematodes, and human, respectively, formed a distinct and well-supported subclade. To exclude the possibility that this clustering was driven primarily by the high conservation of the NBDs, the phylogenetic analysis was repeated using only the TMDs, which represent the most sequence-divergent and substrate-recognition regions of ABC transporters (Sánchez-Fernández et al., 2001). This independent analysis yielded a highly similar tree topology (Fig. 1C), confirming that the observed relationships are not an artefact of NBD conservation.

Together, these *in silico* analyses provide strong evidence that TAP1, NAP8, and ATH12 represent the most compelling candidates for chloroplast-localized ABC peptide exporters in Arabidopsis.

### TAP1, NAP8, and ATH12 localize to the inner membrane of the chloroplast envelope and form homodimers

To experimentally validate the predicted plastid localization of the candidate transporters, *Arabidopsis thaliana* Col-0 plants expressing TAP1, NAP8, or ATH12 C-terminally fused to green fluorescent protein (GFP) were generated and analysed. Chloroplasts isolated from these lines were first sub-fractionated into stromal and membrane-enriched fractions, which were subsequently analysed by immunoblotting to assess GFP distribution (Fig. 2A). In all cases, GFP signals were predominantly detected in the membrane fractions, indicating that the fusion proteins associate with plastid membranes, as expected for transmembrane transporters. Subsequently, protoplasts prepared from the same transgenic lines were examined by confocal laser scanning microscopy (Fig. 2B). For all three transporters, the GFP signal formed a distinct ring surrounding the chlorophyll autofluorescence, a pattern characteristic of chloroplast envelope localization. These observations are consistent with the predicted targeting of TAP1, NAP8, and ATH12 to the plastid envelope membrane system. To further resolve their spatial relationship within the chloroplast envelope, transgenic lines expressing TAP1 fused to GFP and either NAP8 or ATH12 fused to red fluorescent protein (RFP) were generated and crossed to obtain oeTAP1–GFP/oeNAP8–RFP and oeTAP1–GFP/oeATH12–RFP plants. Confocal imaging of mesophyll tissues revealed that both NAP8–RFP and ATH12–RFP displayed ring-shaped fluorescence patterns that fully overlapped with the TAP1–GFP signal (Fig. 2C), indicating co-localization at the same plastid envelope compartment. To specifically mark the inner chloroplast envelope, an oeTAP1–GFP/oeTIC20–RFP line was generated. The TIC20–RFP signal encircled the chlorophyll autofluorescence and overlapped with TAP1–GFP, confirming that TAP1 - and by extension NAP8 and ATH12 - localize to the inner envelope membrane (Fig. 2C). Given that TAP1, NAP8, and ATH12 exhibited indistinguishable subcellular localization patterns and are predicted to function as ABC half-transporters that require dimerization for activity (Kang et al., 2011), their capacity to form homo- or heterodimers was examined using a yeast split-ubiquitin assay (Fig. 2D). Yeast growth on selective medium was observed exclusively in strains expressing identical protein pairs, indicating preferential homodimer formation and no detectable heterodimerization under the conditions tested. Together, these results demonstrate that TAP1, NAP8, and ATH12 co-localize as homodimeric complexes within the inner membrane of the chloroplast envelope, consistent with their proposed function as peptide exporters mediating substrate efflux from the organelle.

**Figure 2.**
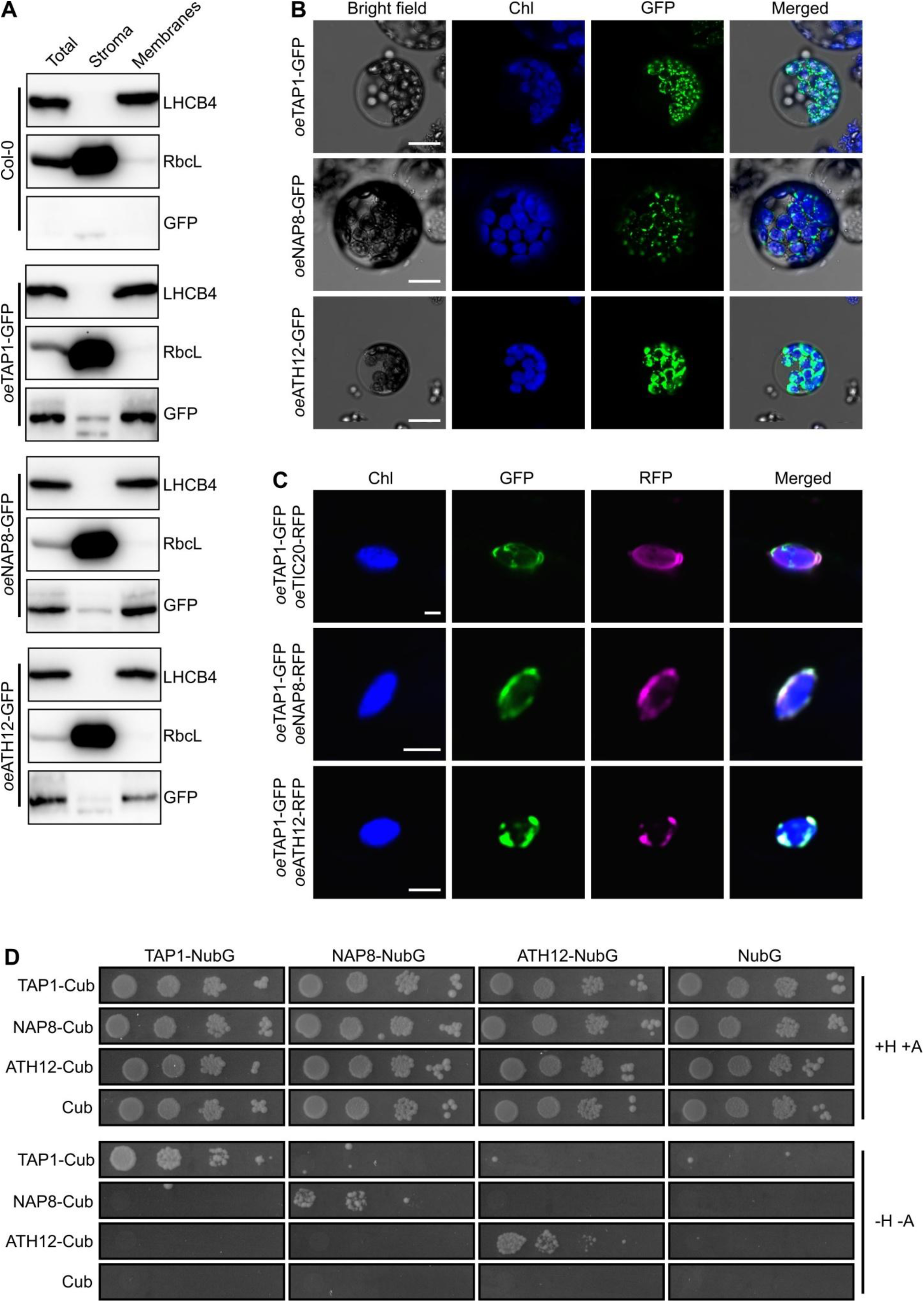
Chloroplast envelope localization and homodimerization of TAP1, NAP8, and ATH12 half-transporters. **A)** Immunoblot analysis of enriched chloroplast fractions from the indicated genotypes. Total chloroplast extracts (Total) were separated into soluble stromal (Stroma) and membrane-enriched (Membranes) fractions. Antibodies against LHCB4 and RbcL were used as markers for membrane and stromal fractions, respectively. GFP antibodies were used to detect the TAP1–GFP, NAP8–GFP, and ATH12–GFP fusion proteins. **B)** Confocal laser scanning microscopy of protoplasts isolated from 18-day-old Col-0 plants overexpressing TAP1–GFP, NAP8–GFP, or ATH12–GFP. Panels show bright-field images, chlorophyll autofluorescence (Chl, blue), GFP fluorescence (green), and merged images. Scale bars, 10 µm. **C)** Confocal images of chloroplasts in mesophyll cells from 18-day-old Col-0 overexpression lines co-expressing TAP1–GFP with either TIC20–RFP, NAP8–RFP, or ATH12–RFP. Chlorophyll autofluorescence (Chl, blue), GFP (green), RFP (magenta), and merged signals are shown. Scale bars, 2.5 µm. **D)** Split-ubiquitin yeast assay to assess protein–protein interactions among TAP1, NAP8, and ATH12. Cub strains expressed the C-terminal half of ubiquitin alone or fused to the indicated transporters, whereas NubG strains expressed the N-terminal half of ubiquitin. Yeast growth on permissive (+H, +A) and selective (–H, –A) media is shown in the upper and lower panels, respectively, indicating homodimerization capacity.

### Arabidopsis TAP1, NAP8, and ATH12 functionally rescue the heat-sensitive phenotype of the yeast *Δmdl1* mutant

In yeast, as noted above, mitochondrial peptide export in response to heat stress is mediated by the ABC transporter Mdl1, and loss of Mdl1 results in a mild but reproducible heat-sensitive phenotype (Young et al., 2001; Jarolim et al., 2013). To assess whether the Arabidopsis ABC transporters TAP1, NAP8, and ATH12 can functionally substitute for Mdl1, we tested their ability to rescue the heat sensitivity of a Δ*mdl1* mutant in *Saccharomyces cerevisiae*. To ensure proper mitochondrial targeting, each plant transporter was expressed in yeast as a fusion with the native Mdl1 mitochondrial targeting peptide. Heat tolerance was assessed using a quantitative liquid growth and recovery assay. Yeast cultures were grown under optimal conditions to exponential phase, split into mock- and heat-treated subcultures, and sampled immediately to generate T_0_ reference points. Mock-treated cultures were maintained at 28 °C, whereas heat-treated cultures were exposed to 45 °C for 2 h. Following treatment, cultures were serially diluted and plated on complete medium and incubated at 28 °C for 3 days to allow colony formation. Cell growth was quantified by calculating the ratio of colony-forming units (CFUs) in heat- or mock-treated samples relative to the corresponding T_0_ reference. Under mock conditions, all strains displayed comparable growth, indicating that expression of the plant transporters did not impair basal yeast fitness (Fig. 3). In contrast, heat-treated cultures exhibited reduced growth across all genotypes, reflecting partial loss of viability during heat exposure. As expected, the Δ*mdl1* mutant showed significantly increased heat sensitivity compared with the wild type. Notably, expression of TAP1, NAP8, or ATH12 in the Δ*mdl1* background fully rescued this phenotype, restoring heat tolerance to levels comparable to those of wild-type cells (Fig. 3). Together, these findings demonstrate that TAP1, NAP8, and ATH12 can functionally replace Mdl1 in yeast. Moreover, the ability of each transporter to independently rescue the Δ*mdl1* phenotype supports their capacity to act as homodimers and indicates functional redundancy among these plant ABC peptide exporters.

**Figure 3.**
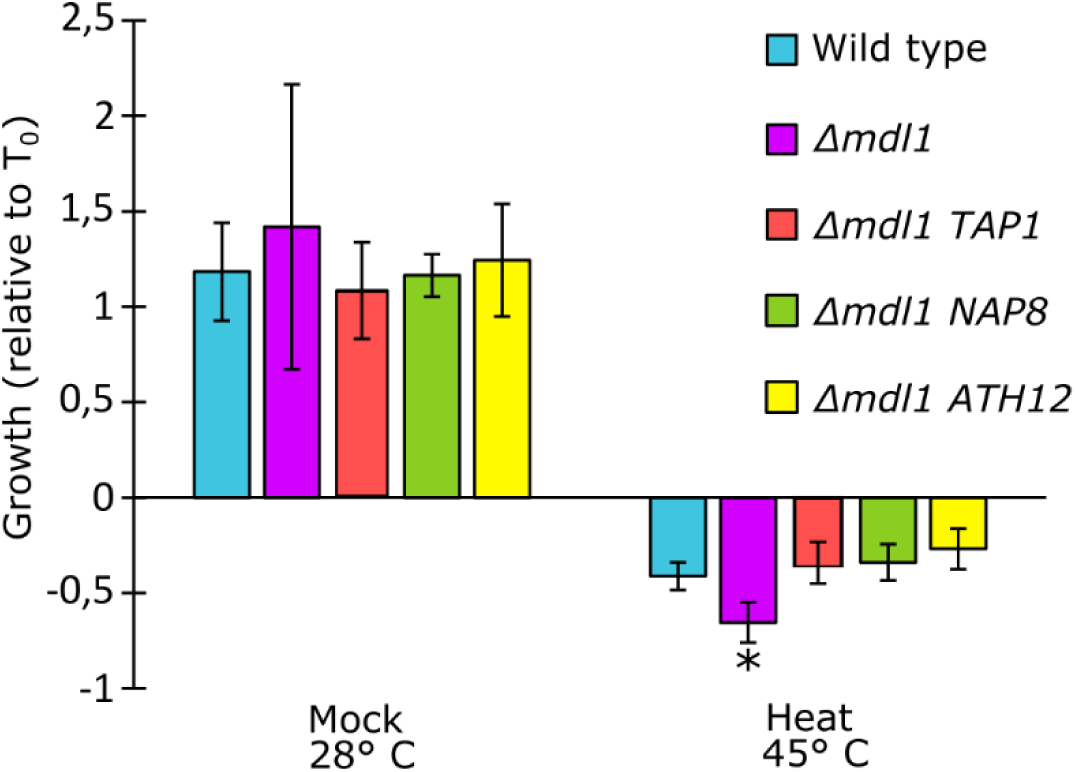
TAP1, NAP8 and ATH12 rescue the heat sensitivity of *mdl1Δ* null mutant. Yeast growth following mock or heat treatment is expressed relative to the corresponding T0 samples for the indicated strains. Asterisks denote statistically significant differences compared with the wild-type heat-treated sample, as determined by Student’s *t*-test (*P* < 0.05). Error bars represent standard deviations from five biological replicates.

### TAP1, NAP8, and ATH12 mediate ATP-dependent peptide export from chloroplasts

To investigate whether TAP1, NAP8, and ATH12 perform functions analogous to those of *Ce*HAF-1 and *Sc*Mdl1, Arabidopsis single mutants carrying T-DNA insertions in the corresponding loci were obtained from the Nottingham Arabidopsis Stock Centre. Putative insertion sites were verified by PCR amplification and sequencing (Fig. S1A), and transcript levels were quantified by qRT-PCR (Fig. S1B). While *TAP1* and *ATH12* expression was completely abolished, *NAP8* transcripts were reduced by approximately 50% relative to the wild type and contained the T-DNA insertion, producing an aberrant transcript consistent with loss of functional expression. The single mutants were subsequently crossed to generate all double-mutant combinations and the corresponding triple mutant. Under optimal growth conditions, none of the mutants displayed overt developmental or morphological phenotypes compared with wild-type plants (Fig. S1C).

To assess peptide export from chloroplasts, we adapted a previously established assay for mitochondria from *Caenorhabditis elegans haf-1* and *Saccharomyces cerevisiae Δmdl1* mutants (Young et al., 2001; Haynes et al., 2010). Intact chloroplasts were isolated by Percoll gradient centrifugation and incubated in ATP-supplemented buffer to support protease and ABC transporter activity. Proteolysis was induced by heat stress at 45 °C for 1 h. After treatment, chloroplasts were gently pelleted, and the supernatants were cleared of protein contaminants by sequential filtration and solid-phase extraction. Peptide content was quantified by UV spectrophotometry. Single and double mutants exhibited peptide release comparable to wild type, whereas the *tap1-1 nap8-1 ath12-1* triple mutant showed a pronounced reduction, releasing approximately half the peptide amount detected in wild-type chloroplasts (Fig. 4A). Chloroplast integrity was confirmed by confocal microscopy, which showed no evidence of envelope rupture or gross structural damage following heat stress, despite partial reduction in chlorophyll autofluorescence (Fig. S2A). Similarly, SDS–PAGE analysis of pre- cleaned supernatants detected only minimal and equivalent amounts of the stromal protein RbcL in wild-type and triple mutant samples (Fig. S2B). Moreover, to assess whether peptide export from chloroplasts is ATP-dependent, the experiment was repeated comparing wild-type and triple mutant chloroplasts, with an additional wild-type sample incubated at 45 °C in the absence of ATP. Notably, peptide levels in the triple mutant supernatant were comparable to those detected in wild-type samples incubated without ATP, whereas peptide release from wild-type chloroplasts approximately doubled in the presence of ATP (Fig. 4B), demonstrating that peptide extrusion is ATP-dependent and that TAP1, NAP8, and ATH12 are functionally redundant.

**Figure 4.**
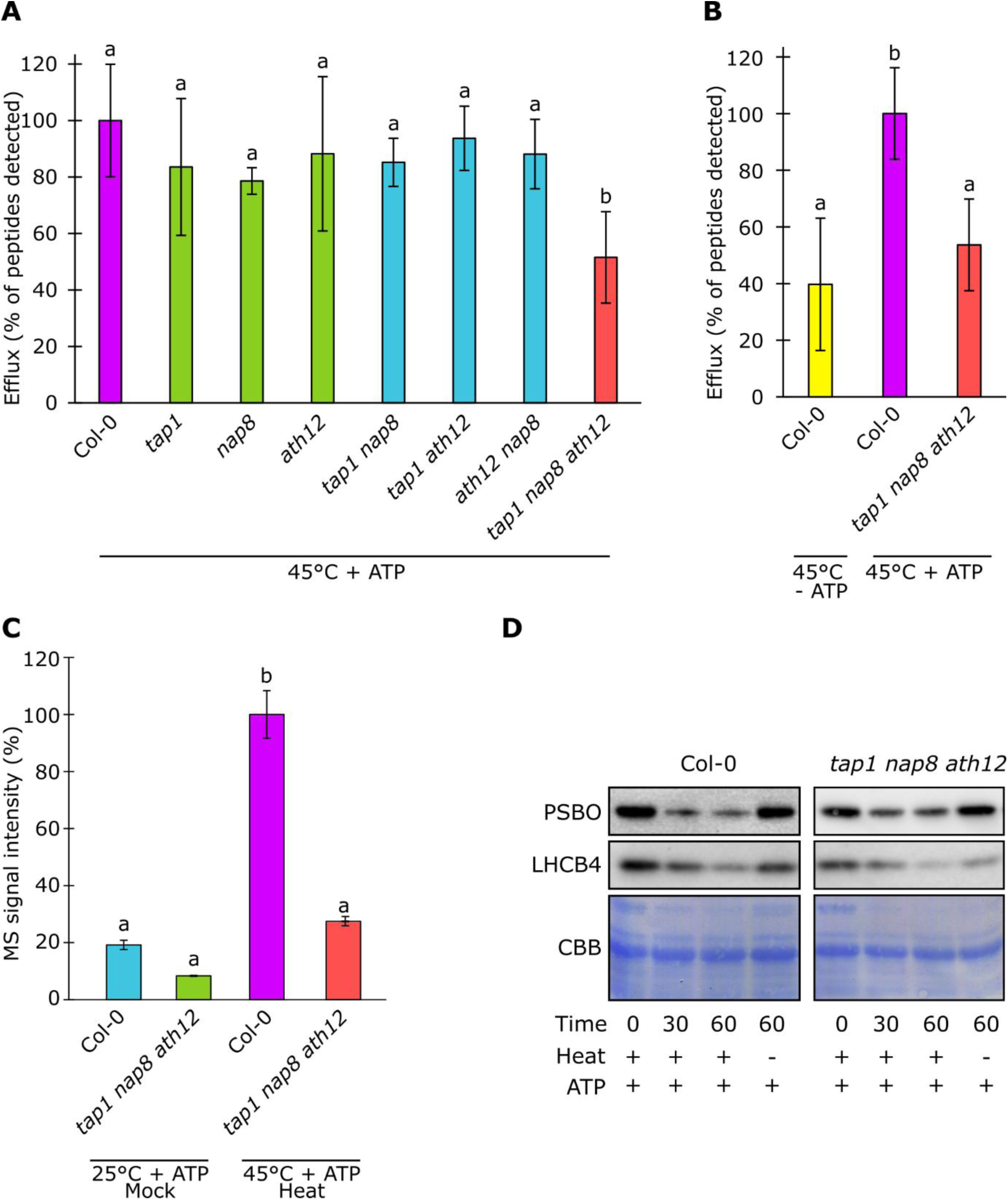
TAP1, NAP8, and ATH12 mediate ATP-dependent peptide export from chloroplasts. **A)** Average normalized peptide efflux from intact chloroplasts isolated from Col-0 (purple), single mutants (green), double mutants (blue), and the triple mutant (red). Peptide efflux was quantified by UV absorbance at 280 nm in the incubation buffer containing ATP after 1 h at 45 °C and normalized to the Col-0 control. Different letters indicate statistically significant differences as determined by one-way ANOVA followed by Tukey–Kramer post hoc test (P < 0.05). Error bars represent standard deviations from at least three independent biological replicates. **B)** Average normalized peptide efflux from chloroplasts incubated for 1 h at 45 °C either in the presence of ATP (Col-0, purple; triple mutant, red) or in the absence of ATP (Col-0, yellow). Values were normalized to the Col-0 + ATP condition. Different letters indicate statistically significant differences (one-way ANOVA with Tukey–Kramer post hoc test, P < 0.05). Error bars represent standard deviations from at least three independent biological replicates. **C)** Average normalized peptide intensities derived from LC–MS/MS analyses of chloroplast supernatants. Values were normalized within each experiment. Different letters indicate statistically significant differences (one-way ANOVA with Tukey–Kramer post hoc test, P < 0.05). Error bars represent standard errors from three independent biological replicates. **D)** Representative immunoblots of chloroplast proteins from wild-type and triple mutant samples incubated under the indicated conditions. Heat (+) indicates incubation for 1 h at 45 °C, whereas Heat (–) corresponds to incubation for 1 h at room temperature. Membranes were probed with antibodies against PSBO and LHCB4. Coomassie Brilliant Blue (CBB) staining is shown as a loading control.

Mass spectrometry was then used to characterize peptides released from chloroplasts. Wild-type and triple mutant chloroplasts were incubated under mock (room temperature) or heat stress conditions (45 °C) in the presence of ATP. Total peptide abundance, normalized across three biological replicates, increased significantly only in heat-treated wild-type samples, whereas the triple mutant showed no comparable increase (Fig. 4C, Table S2), confirming impaired peptide export in the absence of the three transporters. Analysis of peptide origin revealed that the most abundant peptides derived predominantly from thylakoid membrane proteins, including multiple components of the photosystem II oxygen-evolving complex (PSBO1, PSBO2, PSBQ1, PSBQ2, PSBP1, PSBR) and several light-harvesting complex proteins (LHCB1, LHCB4, LHCB5) (Tables S2–S3).

To exclude the possibility that reduced peptide efflux in the triple mutant was due to altered proteolysis, chloroplasts were analysed by immunoblotting for PSBO and LHCB4 during heat stress. Protein levels decreased similarly in wild-type and triple mutant chloroplasts over the 60-min incubation at 45 °C, whereas incubation at room temperature with ATP did not affect protein stability (Fig. 4D). These results indicate that the diminished peptide release observed in triple mutant chloroplasts reflects the absence of TAP1, NAP8, and ATH12, rather than differences in proteolytic activity.

### Heat stress increases the abundance of thylakoid-derived peptides in Arabidopsis chloroplasts

Overall, LC–MS/MS analyses across all experimental conditions led to the identification of 15,016 non-redundant peptide sequences (Table S2), originating from 562 source proteins, each contributing more than 2 peptides (Table S3). For all downstream analyses, only peptides consistently detected in all three biological replicates per condition were retained. Under mock conditions, 1,672 peptides were detected in Col-0, whereas 1,263 peptides in *tap1 nap8 ath12* samples. Following heat stress, the number of non-redundant peptides increased to 2,558 in Col-0 and 1,973 in the triple mutant. The consistently lower number of peptides detected in the triple mutant under both mock and heat conditions further supports a reduced chloroplast peptide efflux in the absence of TAP1, NAP8, and ATH12, in agreement with the transport assays (Fig. 4). Identified peptides ranged in length from 5 to 25 amino acids, with an average length of 15–16 residues (Table S2; Fig. 5A). Peptide length distributions were comparable across all genotypes and treatments (Fig. S2C). However, when peptide abundances were normalized to the total MS signal intensity of Col-0 samples, a marked reduction in peptide abundance was observed in the triple mutant under both mock and heat conditions (Fig. 5A), further corroborating impaired peptide export. Source proteins contributing peptides under all tested conditions were ranked according to the number of detected peptides (Table S4). Only proteins identified by at least 1 peptide in 2 out of 3 biological replicates were considered. In all samples, more than 97% of peptides mapped to plastid-localized proteins, indicating comparable purity of the isolated chloroplast preparations (Table S4; Fig. S2D). Plastid-derived peptides were subsequently grouped based on the sub-plastid localization of their source proteins (Fig. 5B; Table S4). In heat-treated Col-0 samples, peptides predominantly originated from thylakoid-associated proteins, including integral, peripheral, and lumenal proteins, accounting for over 80% of the total peptide pool. Stromal proteins contributed approximately 11%, while envelope- and plastoglobule-derived peptides accounted for about 2% and <1%, respectively. Under mock conditions, the relative contribution of thylakoid-derived peptides decreased to approximately 70%, accompanied by an increased contribution from stromal proteins (∼16%), consistent with enhanced turnover of thylakoid proteins during heat stress. Importantly, the overall sub-plastid distribution of peptides in the triple mutant closely mirrored that observed in Col-0 (Fig. 5B; Table S4).

**Figure 5.**
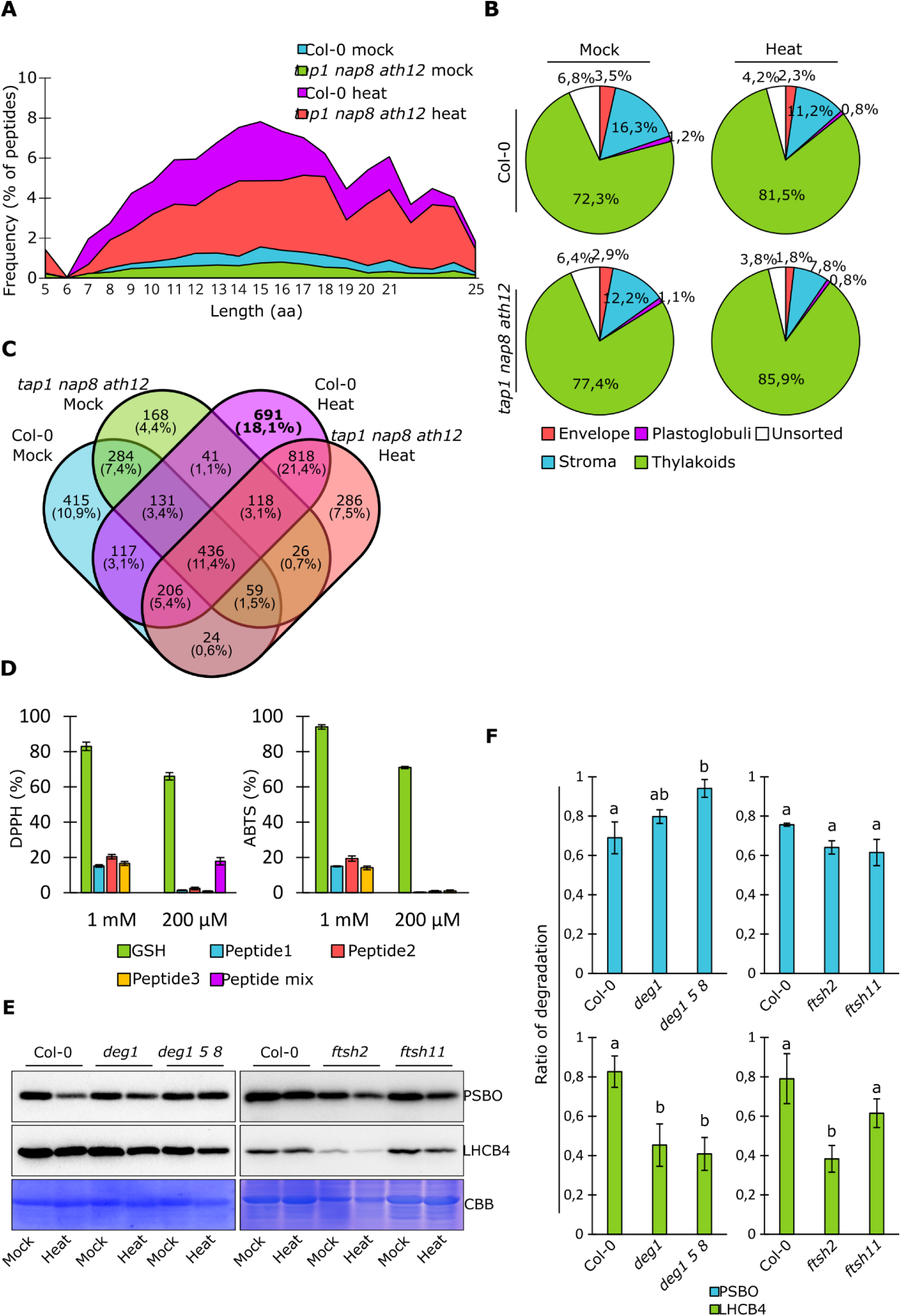
Molecular characterization of chloroplast-exported peptides and their functional properties. **A)** Frequency distribution of peptide lengths (in amino acids) detected in the incubation buffer of chloroplasts exposed to 45 °C in the presence of ATP, based on LC–MS/MS analyses from at least three independent experiments. Data were normalized to the total MS signal intensity of heat-treated Col-0 samples. **B)** Relative abundance of peptides detected in the incubation buffer of wild-type and *tap1 nap8 ath12* chloroplasts under control conditions or following heat treatment (45 °C, ATP present), grouped according to the sub-plastid localization of their source proteins. Percent contributions of each compartment are indicated. Data derive from three independent LC–MS/MS experiments. **C)** Venn diagram illustrating the overlap of non-redundant peptide sequences detected across samples. Peptides uniquely induced by heat treatment in Col-0 are highlighted in bold. **D)** In vitro antioxidant activity of Peptide1, Peptide2, Peptide3, and their mixture, assessed by DPPH and ABTS radical scavenging assays. Glutathione (GSH) was included as a reference compound. Error bars represent standard deviations from at least three independent replicates. **E)** Representative immunoblots of total protein extracts from cycloheximide-treated leaves of the indicated genotypes incubated at 25 °C (Mock) or 45 °C (Heat) for 2 h. Membranes were probed with antibodies against PSBO and LHCB4. Coomassie Brilliant Blue (CBB)-stained gels are shown as loading controls. **F)** Quantification of protein degradation, expressed as the ratio between Heat and Mock signal intensities for the indicated genotypes and target proteins. Different letters indicate statistically significant differences as determined by one-way ANOVA followed by Tukey–Kramer post hoc test (*P* < 0.05). Error bars represent standard errors from at least three independent biological replicates.

Visualization of non-redundant peptide sequences using Venn diagrams revealed complex overlap patterns among samples (Fig. 5C; Table S2). Approximately 20% of peptides (691 sequences derived from 176 proteins) were uniquely detected in heat-treated Col-0 samples. Nevertheless, 818 peptides were shared between heat-treated Col-0 and triple mutant samples, indicating the release of a common subset of peptides in response to heat stress. Physicochemical characterization of Col-0-specific heat-induced peptides showed that 48% carried a net negative charge (–7 to –1), 29% were positively charged (+1 to +3), and 23% were neutral (Fig. S2E; Table S5). The weighted average normalized hydrophobicity of these peptides was –0.34 (Moon and Fleming scale), indicating that they are predominantly hydrophilic and unlikely to cross biological membranes by passive diffusion (Fig. S2F; Table S5). Functional prediction analyses further revealed that only 6.8% of heat-induced peptides were classified as antimicrobial, whereas the vast majority (∼95%) were predicted to possess antioxidant activity (Table S5). Notably, comparison of these peptides with the 270 peptides released from yeast mitochondria in an ATP- and temperature-dependent manner (Augustin et al., 2005) revealed striking similarities in multiple physicochemical parameters, including average linear moment, normalized hydrophobic moment, predicted membrane penetration depth, and propensity to adopt a polyproline II (PPII) conformation (Table S6). Both datasets also displayed comparable functional predictions, with approximately 5.0% of mitochondrial peptides predicted to be antimicrobial and about 96% predicted to exhibit antioxidant activity.

The potential antioxidant activity of exported chloroplast peptides was further assessed *in vitro*. Three 12 aa long peptides were synthesized. Peptide1 (SVSKNAPPEFQN) and Peptide2 (SDTDLGAKVPKD), corresponding to shared sequences derived from PSBO1-originated peptides, and Peptide3 (SLDQNLAKNLAG), derived from LHCB4-originated peptides (Table S5). Radical scavenging activity was evaluated using DPPH and ABTS assays at high (1 mM) and low (200 µM) concentrations, with glutathione (GSH) serving as a reference (Fig. 5D). At high concentrations, individual peptides exhibited 15-20% scavenging activities in both DPPH and ABTS assays, compared with 80-90% observed for GSH. At lower concentrations, peptide activity was negligible, whereas GSH retained ∼70% activity. Notably, a mixture of the three peptides displayed ∼18% DPPH scavenging activity, suggesting that the exported peptide pool may act synergistically. Together, these results support the notion that chloroplast-exported peptides possess antioxidant potential.

Among the 176 proteins uniquely contributing peptides to heat-treated Col-0 samples, the most prominent sources were the 33-kDa subunit of the PSII oxygen-evolving complex (PSBO1) and the 16-kDa subunit (PSBQ2), which generated 54 and 38 peptides, respectively. Within the light-harvesting complex, the chlorophyll a/b-binding protein CP29.2 (LHCB4.2) was the major contributor, yielding 17 peptides. Mapping of peptide sequences revealed a strong enrichment in the C-terminal regions of oxygen-evolving enhancer proteins (residues 87–328 for PSBO1 and 88–219 for PSBQ2), as well as in regions spanning residues 45–65 and 90–120 of LHCB4 (Fig. S3). Conserved core peptide motifs were identified within peptides derived from PSBO1, PSBQ2, and LHCB4 (Table S7). In PSBO1-derived peptides, recurrent core sequences included PPEFQ and GKPDS (each detected in nine peptides) and DTDLGAKVPK (detected in eight peptides). In contrast, the core sequence AKPKE was most abundant among PSBQ2-derived peptides, appearing in 12 sequences, followed by SLKDL and NLDYAAR, each detected in 10 peptides. To gain insights into the proteolytic activities responsible for peptide generation during heat stress, cleavage motif analysis was performed by aligning amino acid sequences spanning ±15 residues around the N- and C-terminal cleavage sites of Col-0 specific heat-induced peptides (Fig. S2G). At N-terminal cleavage sites, a strong preference for alanine, leucine, arginine, and lysine was observed at the P1 position, whereas no clear preference emerged at P–1. Upstream regions (positions –1 to –14) were enriched in basic residues, particularly lysine and arginine, as well as alanine and leucine. At C-terminal cleavage sites, arginine, lysine, glutamic acid, and leucine were strongly enriched at P1’, again with no pronounced P–1’ preference. Given that DEG and FTSH proteases play central roles in chloroplast protein quality control under stress conditions (see Introduction), and that their reported cleavage specificities are consistent with the identified motifs, their contribution to heat-induced peptide generation was examined using protease-deficient *Arabidopsis* mutants. Specifically, mutants lacking luminal proteases involved in PSII repair and protein quality control (*deg1* and *deg1 deg5 deg8*), as well as mutants deficient in the thylakoid membrane protease FTSH2 and the inner-envelope protease FTSH11, were analysed. Leaves were pretreated with cycloheximide to inhibit cytosolic protein synthesis and then incubated at 45 °C for 2 h or maintained at room temperature as controls. Immunoblot analyses revealed reduced PSBO degradation in *deg* mutants relative to wild-type plants, with PSBO levels remaining unchanged in both mock- and heat-treated *deg1 deg5 deg8* samples. In contrast, LHCB4 degradation was enhanced in *deg* mutants. Conversely, *ftsh2* mutants exhibited increased degradation of both PSBO and LHCB4, whereas *ftsh11* mutants behaved similarly to wild-type plants (Fig. 5 E, F).

### The concomitant loss of TAP1, NAP8 and ATH12 Impairs Heat Tolerance and Redox Homeostasis in Arabidopsis

To determine whether the Arabidopsis *tap1 nap8 ath12* triple mutant is sensitive to high temperature, similar to *C. elegans haf-1* and *S. cerevisiae Δmdl1* mutants, plantlets grown in sterile dishes for 15 days after sowing (DAS) were exposed to 45°C for 2 hours. After heat treatment, plantlets were returned to optimal growth conditions for 2 days before analysis. As controls, Col-0 and the triple mutant were grown under optimal conditions without heat stress. Treated triple mutant plantlets appeared paler and smaller than wild-type plants, indicating increased heat sensitivity (Fig. 6A). Consistently, biomass accumulation in the mutant was significantly reduced compared to the wild type (Fig. 6B). Furthermore, quantification of chlorophylls and carotenoids revealed a marked decrease in the mutant (Fig. 6C–E). Given the heightened heat sensitivity of the triple mutant, CLPB3 accumulation was monitored under control conditions, after 2 h at 45°C, and following 1 and 3 h of recovery. CLPB3 accumulation was markedly altered in the mutant, showing higher levels than Col-0 under control conditions, followed by a progressive decrease after heat stress and throughout recovery. In contrast, wild-type plants exhibited the opposite trend, with a gradual increase in CLPB3 accumulation in response to heat stress and during recovery (Fig. 6F, G). Notably, CLPB3 transcript levels were similarly upregulated in both wild-type and mutant plantlets upon heat treatment (Fig. 6H), indicating that the observed differences in protein accumulation arise from post-transcriptional regulation.

**Figure 6.**
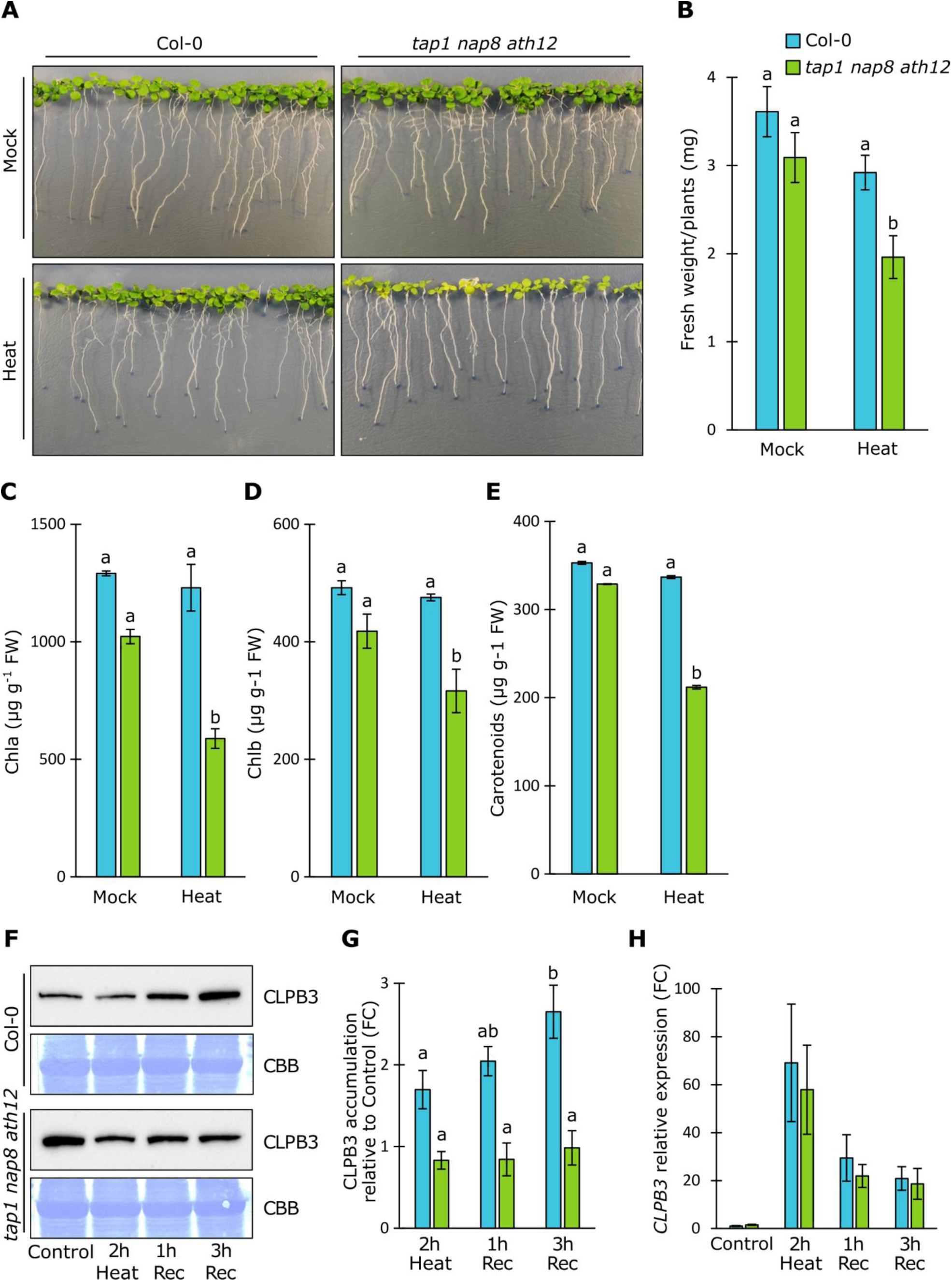
Characterization of the heat-sensitive phenotype of the *tap1 nap8 ath12* triple mutant. **A)** Representative images of 15-day-after-sowing (DAS) plantlets incubated either under control conditions or at 45 °C for 2 h. **B)** Fresh weight of plantlets shown in (A). Different letters indicate statistically significant differences as determined by one-way ANOVA followed by Tukey–Kramer post hoc test (*P* < 0.05). Error bars represent standard errors from at least three independent biological replicates. **C–E)** Quantification of chlorophyll a (C), chlorophyll b (D), and total carotenoids (E) of plantlets shown in (A). Different letters indicate statistically significant differences (one-way ANOVA with Tukey–Kramer post hoc test, *P* < 0.05). Error bars represent standard errors from at least three independent biological replicates. **F)** Representative immunoblots of total protein extracts from wild-type and triple mutant plantlets collected before heat treatment, after 2 h at 45 °C, and following 1 h and 3 h of recovery under optimal conditions. Membranes were probed with antibodies against CLPB3. Coomassie Brilliant Blue (CBB) staining is shown as a loading control. **G)** Quantification of CLPB3 protein abundance from immunoblot analyses shown in (G). Different letters indicate statistically significant differences (one-way ANOVA with Tukey–Kramer post hoc test, *P* < 0.05). Error bars represent standard errors from 10 independent biological replicates. **H)** Representative relative CLPB3 transcript levels before heat treatment, after 2 h at 45 °C, and following 1 h and 3 h of recovery under optimal conditions. Error bars represent standard errors.

The increased heat sensitivity of the triple mutant, together with the identification of heat-induced peptides uniquely detected in wild-type samples and predicted to possess antioxidant activity (Table S5; Fig. 5D), prompted us to investigate alterations in cellular redox homeostasis. Redox parameters were therefore analyzed before heat stress, during heat treatment, and after 3 h of recovery. Protein oxidation and lipid peroxidation were assessed as indicators of oxidative stress. In the triple mutant, protein oxidation levels remained largely unchanged across all time points, whereas wild-type samples exhibited a significant increase upon heating, followed by a return to basal levels during recovery (Fig. 7A). In contrast, lipid peroxidation increased markedly in the mutant during heat stress but reverted to baseline values during recovery, while no major changes were observed in wild-type plants (Fig. 7B). We next quantified the cellular pools of ascorbate (ASC) and glutathione (GSH), two major hydrophilic antioxidants. In the triple mutant, both ASC and GSH accumulated already during heat stress and remained elevated throughout recovery. In wild-type plants, however, increased levels of these antioxidants were mainly observed during the recovery phase (Fig. 7C, D). Notably, the basal GSH content was lower in the mutant than in the wild type, and even at its maximum accumulation during heating, GSH levels in the mutant did not reach those observed in Col-0 after 3 h of recovery (Fig. 7D). To further characterize the antioxidant capacity of the mutant, the activity and expression of key ROS-scavenging enzymes—superoxide dismutases (SODs), catalases (CATs), and ascorbate peroxidases (APXs)—were analysed. Basal SOD activity was lower in the triple mutant than in the wild type. In the mutant, SOD activity peaked during heat treatment and subsequently returned to basal levels during recovery, whereas in wild-type plants the highest activity was observed during recovery (Fig. 7E). APX activity was also overall reduced in the mutant compared with the wild type; however, in the mutant it progressively increased during both heat treatment and recovery, while wild-type plants showed a transient inhibition during heating followed by recovery at later stages (Fig. 7F). CAT activity remained relatively stable in both genotypes, although a reduction relative to basal levels was observed in wild-type plants during heat stress (Fig. 7G).

**Figure 7.**
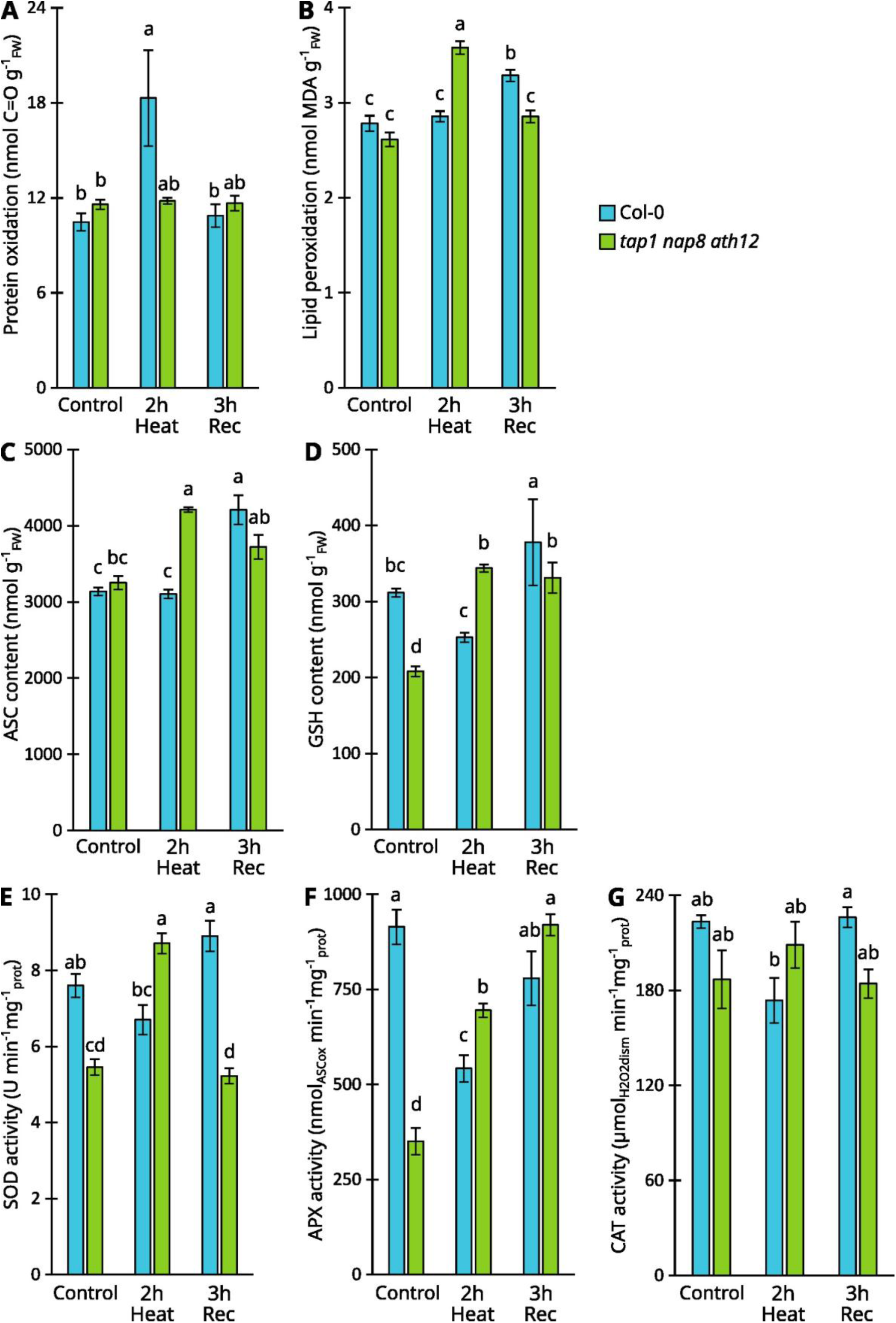
Altered redox homeostasis in response to heat stress. **A)** Levels of oxidized proteins measured in plantlets before heat treatment, immediately after 2 h incubation at 45 °C, and following 3 h of recovery under optimal conditions. **B)** Lipid peroxidation levels measured in plantlets before heat treatment, immediately after 2 h at 45 °C, and after 3 h of recovery. **C)** Ascorbate (ASC) content measured in plantlets before heat treatment, immediately after 2 h at 45 °C, and following 3 h of recovery. **D)** Glutathione (GSH) content measured in plantlets before heat treatment, immediately after 2 h at 45 °C, and following 3 h of recovery. **E)** Superoxide dismutase (SOD) activity measured in plantlets before heat treatment, immediately after 2 h at 45 °C, and after 3 h of recovery. **F)** Ascorbate peroxidase (APX) activity measured in plantlets before heat treatment, immediately after 2 h at 45 °C, and following 3 h of recovery. **G)** Catalase (CAT) activity measured in plantlets before heat treatment, immediately after 2 h at 45 °C, and after 3 h of recovery. For all panels, different letters indicate statistically significant differences as determined by one-way ANOVA followed by Tukey–Kramer post hoc test (*P* < 0.05). Error bars represent standard errors from at least three independent biological replicates.

These enzymatic trends were mirrored by distinct transcriptional responses of the corresponding antioxidant genes (Fig. S4). Specifically, *Cu/ZnSOD1*, *Cu/ZnSOD2*, and *FSD3*, which displayed reduced transcript levels in the mutant under control conditions, were strongly induced upon heat treatment in the mutant but repressed in the wild type (Fig. S4A, B, E). *FSD1* expression increased in both genotypes during recovery (Fig. S4C), whereas *FSD2* failed to be induced during recovery in the mutant (Fig. S4D). Moreover, *APX1*, *APX2*, and *tAPX* exhibited a stronger transcriptional response to heat stress in the mutant than in wild-type plants (Fig. S4F–H). Similarly, *CAT2* expression was higher in the mutant during heating compared with Col-0 (Fig. S4I).

## Discussion

Protein quality control is essential for chloroplast function, particularly under heat stress, which imposes severe challenges on photosynthetic membranes and proteostasis. In this study, we identify TAP1, NAP8, and ATH12 as plastid-localized ABCB half-transporters that collectively mediate ATP-dependent peptide export from chloroplasts and are required for effective heat stress tolerance in *Arabidopsis thaliana*. Our findings uncover a previously unrecognized chloroplast peptide export pathway that functionally parallels mitochondrial peptide extrusion systems described in yeast and metazoans, while revealing a plant-specific adaptive response to thermal stress.

### TAP1, NAP8, and ATH12 function as plastid peptide exporters

Independent evidence supports the role of TAP1, NAP8 and ATH12 as plastid peptide exporters. First, *in silico* analyses combined with phylogenetic reconstruction place TAP1, NAP8, and ATH12 firmly within the ABCB half-transporter family and reveal a close evolutionary relationship with well-characterized peptide exporters, including *Sc*Mdl1, *Ce*HAF-1, and mammalian TAP/TAPL proteins (Fig. 1; Young et al., 2001; Sánchez-Fernández et al., 2001; Wolters et al., 2005; Haynes et al., 2010; Kang et al., 2011; Lehnert et al., 2016; Nöll et al., 2017). Notably, this clustering persists when analyses are restricted to the transmembrane domains, highly divergent portions and responsible for substrate recognition, indicating that functional relatedness extends beyond the highly conserved nucleotide-binding domains (Sánchez-Fernández et al., 2001; Stieger and Higgins, 2007; Kang et al., 2011). Together, these data strongly support the notion that TAP1, NAP8, and ATH12 represent bona fide plastid homologs of mitochondrial peptide extruders.

Second, the ability of TAP1, NAP8, and ATH12 to fully rescue the heat-sensitive phenotype of the yeast *Δmdl1* mutant provides compelling evidence for deep functional conservation of peptide export mechanisms (Fig. 3). In yeast, Mdl1 mediates the ATP-dependent extrusion of peptides generated by mitochondrial proteolysis, thereby contributing to organellar proteostasis (Young et al., 2001; Augustin et al., 2005; Jarolim et al., 2013). Functional substitution by TAP1, NAP8, or ATH12 indicates that both transport mechanisms and substrate properties are evolutionarily conserved, despite the distinct proteomes and stress sensitivities of mitochondria and chloroplasts. This observation further supports the idea that substrate recognition is not strictly sequence-dependent but instead relies on shared physicochemical features. In chloroplasts, where heat stress disproportionately affects thylakoid membrane complexes (Allakhverdiev et al., 2008), an efficient peptide export system may be essential to prevent accumulation of potentially bioactive degradation products and to maintain cellular homeostasis. (Fig. 5D, Table S5; Ferro et al., 2014).

Third, biochemical and mass spectrometry analyses demonstrate that heat stress triggers a pronounced increase in ATP-dependent peptide export from wild-type chloroplasts, whereas this response is strongly attenuated in the *tap1 nap8 ath12* triple mutant (Fig. 4 and 5, Table S2). The exported peptides predominantly originate from thylakoid-associated proteins, including components of the photosystem II oxygen-evolving complex and light-harvesting antennae - well-established targets of heat-induced damage (Fig. 5B, Tables S3 and S4; Allakhverdiev et al., 2008). This pattern closely mirrors observations in mitochondria, where exported peptides mainly derive from respiratory chain components (Young et al., 2001; Augustin et al., 2005; Haynes et al., 2010). Plastid extruded peptides displayed a length distribution ranging from 5 to 25 residues, with the most abundant being of 15-16 amino acids long (Fig. 5A, Fig. S2C). These data are in good agreement with results from previous studies across different organisms and organelles, thereby supporting our findings. In fact, peptides extruded from *C. elegans* and *S. cerevisiae* mitochondria ranged from 6 to 30 amino acids in length, with a median length of 14 amino acids (Augustin et al., 2005; Haynes et al., 2010). Most of the Col-0 heat-induced peptides carried a net negative charge (ranging from -7 to -1 net charges; Fig. S2E). Noteworthy, the negative weighted average normalized hydrophobicity of these peptides indicated that the peptides were predominantly hydrophilic and highly soluble in water, making passive diffusion across a biological membrane unlikely (Fig. S2F). Only a small fraction (6.8%) of these HS-induced peptides was predicted to be antimicrobial, with the majority (95%) instead predicted to be antioxidant (Table S5). Peptides released from *S. cerevisiae* mitochondria in an ATP-and temperature-dependent manner (Augustin et al., 2005) exhibited comparable physicochemical properties, including predicted antimicrobial and antioxidant activities (Table S6).

Finally, experimental localization analyses place TAP1, NAP8, and ATH12 at the inner envelope membrane of chloroplasts (Fig. 2), positioning them ideally to mediate export of stromal or thylakoid-derived peptides to the cytosol (Inaba and Schnell, 2008). This result is consistent with a previous study of chloroplast fractions analysed by mass spectrometry (Ferro et al., 2010). Their exclusive homodimerization (Fig. 2D) further supports functional independence, a feature commonly observed among ABC half-transporters, including HAF-1 and Mdl1 (Schaedler et al., 2015). Additionally, TAP1, NAP8 and ATH12 display functional redundancy, as the peptide export from chloroplasts is hampered only if all three transporters are absent (Fig. 4A and 5A). This redundancy likely accounts for the lack of overt phenotypes in single and double mutants under optimal and stress conditions (Fig. S1C) and for the ability of each transporter to rescue the heat-sensitive phenotype of the yeast *Δmdl1* mutant (Fig. 3).

### Heat-induced peptide export reflects thylakoid protein turnover

The heat-induced peptides uniquely released from Col-0 chloroplasts predominantly originated from thylakoid-associated proteins involved in photosystem II (PSII) structure and function, as well as in light harvesting (Fig. 5C, Table S5). Notably, many of these proteins, including D1, D2, CP43, LHCB4, and the oxygen-evolving complex (OEC) subunits, are well-established substrates of proteolytic degradation during photodamage and heat stress, a process primarily mediated by DEG and FTSH proteases (Huesgen et al., 2009; Kato and Sakamoto, 2018). Our analysis revealed that the degradation kinetics of representative thylakoid proteins, such as PSBO and LHCB4, were comparable between wild-type and triple mutant chloroplasts, indicating that heat-induced protein degradation per se is not impaired in the mutant background (Fig. 4D). Given that PSBO1 is a nuclear-encoded thylakoid protein associated with the lumenal surface of PSII (De Las Rivas and Barber, 2004; Roose et al., 2007), its proteolysis is most likely mediated by proteases localized in the thylakoid lumen. Consistently, our cleavage motif analysis, together with the strong inhibition of PSBO degradation observed in the *deg1 deg5 deg8* triple mutant, supports the involvement - at least in part - of DEG proteases in the generation of heat-induced peptides (Fig. S2G, Fig. 5E, F). DEG proteases are ATP-independent enzymes that are strongly activated under stress conditions, including high light and elevated temperature (Nishimura et al., 2017; Luciński and Adamiec, 2023). Chloroplast DEG proteases do not have a single strict cleavage residue, but they show a clear preference for hydrophobic amino acids (Krojer et al., 2008). The observed frequencies of amino acids in the P1 position indicated a strong preference for small hydrophobic residues, including alanine which was among the most overrepresented residues in our cleavage site analysis (Fig. S2G). Moreover, consistent with the observed cleavage patterns, several chloroplast-localized Deg/HtrA proteases, such as DEG1, DEG5, and DEG8 contain trypsin-like domains capable of cleaving after basic residues, including arginine and lysine (Schuhmann et al., 2008; Sun et al., 2010; Chen et al., 2014). In contrast, immunoblot analyses revealed no reduction in LHCB4 degradation in the *deg* mutant lines (Fig. 5E, F), indicating that protease families other than DEG also contribute to peptide generation under heat stress. Mapping of LHCB4-derived peptides showed that they originate from the stromal side of the protein (Table S7, Fig S3), suggesting the involvement of stromal proteases in their production. The major ATP-dependent protease families localized in the chloroplast stroma - CLP, LON1, and FTSH - are well known for their roles in heat stress tolerance and in the removal of misfolded or oxidized proteins (Kapri-Pardes et al., 2007; Nishimura et al., 2017; Luciński and Adamiec, 2023). However, immunoblot analyses of *ftsh2* and *ftsh11* mutant did not reveal any reduction in the degradation of either PSBO or LHCB4, thereby excluding a major contribution of this protease to the generation of heat-induced peptides from these substrates (Fig. 5E, F).

Once exported from chloroplasts, peptides may undergo different cellular fates and possess bioactive properties (Ferro et al., 2014). Indeed, a notable feature of heat-induced peptides uniquely detected in wild-type samples is their potential antioxidant activity as observed both *in silico* and *in vitro* (Fig. 5D; Table S5). Importantly, the antioxidant properties of a mixture of 3 peptides appeared synergistically enhanced *in vitro* with respect to single peptides. These observations, while not conclusive, support that exported peptides might have redox-buffering potential.

Furthermore, the presence of shared core sequence motifs among these peptides (see Table S7, Fig. S3) suggests that they are unlikely to represent random degradation products. Rather, the conservation of these core regions points to selective protection from complete chloroplast proteolysis and may facilitate recognition by specific cytosolic ligands, thereby enabling downstream signalling events. This non-random degradation pattern is reminiscent of that described by Lyapina and colleagues (Lyapina et al., 2021) and is consistent with earlier observations by Augustin et al. (2005), who reported similar features for peptides exported by the mitochondrial Mdl1 transporter.

Although the functional significance of these conserved peptide sequences remains unclear, accumulating evidence indicates that peptides can play critical roles in the regulation of gene expression, particularly under stress conditions. In *Arabidopsis thaliana*, exposure to high-light stress leads to the sensing of elevated singlet oxygen (^1^O₂) levels by the EX1 protein via the Trp643 residue within its DUF/SOS domain. This oxidative signal triggers EX1 cleavage by the FTSH2 protease, resulting in the release of a defined peptide corresponding to the UVR domain (DRLLSVLKSQLNRAIKREDYEDAARLK; amino acids 127–153). This peptide is subsequently exported from the chloroplast to the cytosol through a still-unknown mechanism involving the TOC33 protein. Once in the cytosol, the EX1-UVR-derived peptide functions as a mobile retrograde signal, translocating to the nucleus where it modulates the expression of singlet-oxygen-responsive genes through interactions with transcription factors such as WRKY18 and WRKY40 (Zhao et al., 2025). Evidence for DNA–peptide interactions have also been reported in humans, where peptides have been shown to influence DNA methylation status (Khavinson et al., 2021). In stem cells, for example, the short peptide AEDG epigenetically regulates gene expression and protein synthesis through direct interactions with histones (Khavinson et al., 2020).

Together, these findings raise the possibility that chloroplast-derived peptides contribute to cytosolic redox balance and/or stress signalling through gene expression regulation, a hypothesis that warrants future investigation.

### Impaired peptide export compromises redox homeostasis and heat tolerance

The *tap1 nap8 ath12* triple mutant displays pronounced heat sensitivity at the whole-plant level, accompanied by altered redox dynamics (Fig. 6, Fig. 7 and Fig. S4). Immediately after heat stress, ASC/GSH levels and the activities of detoxifying enzymes rise, indicating a reinforced early redox protective response. Considering that plant-derived peptide can display intrinsic antioxidant activity (César et al., 2024; Zhu et al., 2024), supported by the radical scavenging capacity measured for chloroplast-origin peptides in this study (Fig. 5D), a possible explanation is that the increase in antioxidants represents a compensatory but ultimately insufficient adjustment of redox balance. This pattern resembles that of other mutants with impaired chloroplast quality control, in which perturbed proteostasis or retrograde signalling coexists with heightened antioxidant activity but persistent stress sensitivity (Schwenkert et al., 2022; Fu et al., 2022; Wang et al., 2025). In this view, reinforcement of enzymatic defences would be expected to dampen the short ROS transient that normally contributes to initiating heat-stress signalling and acclimation (Sun and Guo, 2016; Li and Kim, 2022; Foyer and Hanke, 2022; Lasorella et al., 2022; Fortunato et al., 2023). Notably, during the recovery phase, despite sustained high antioxidant activity and the induction of most antioxidant transcripts, SOD activity and FSD2 transcript abundance remain lower in the triple mutant than in the wild type. FSD2, also known as PAP9, functions together with FSD3 at thylakoids and plastid nucleoids to detoxify superoxide and support photosynthetic performance (Myouga et al., 2008; Gallie and Chen, 2019). PAP9/FSD2 associates with the plastid-encoded RNA polymerase (PEP), helping to stabilize and modulate the complex and protecting PEP under heat oxidative conditions (Favier et al., 2021; Vergara-Cruces et al., 2024; Chen et al., 2025). Together, the reduced FSD2/PAP9 transcript levels and diminished SOD activity indicate persistent thylakoid oxidative constraints and less efficient PEP-dependent transcription, factors expected to contribute to the observed thermosensitivity. Moreover, CLPB3 altered protein accumulation, despite its normal transcriptional induction, points to post-transcriptional limitations, potentially influenced by the altered redox homeostasis. Since CLPB3 is a key chloroplast chaperone required for resolution of heat-induced protein aggregates and for sustaining thermotolerance (Parcerisa et al., 2020; Kreis et al., 2023), its impaired accumulation could contribute to exacerbate the heat-sensitive phenotype of the triple mutant.

## Conclusions

Collectively, our data support a model in which TAP1, NAP8, and ATH12 constitute a chloroplast peptide export system that alleviates heat-induced proteotoxic stress by removing degradation products generated primarily from thylakoid proteins. Loss of this pathway results in mis-timed redox responses, impaired accumulation of stress-adaptive proteins, and reduced thermotolerance. This study extends the concept of organellar peptide export beyond mitochondria and identifies chloroplast peptide efflux as a critical, previously overlooked component of plant heat stress resilience. Understanding the downstream fate and signalling role of exported peptides will be an important next step toward elucidating how chloroplasts communicate stress status to the nucleus and the rest of the cell.

## Material and methods

### Identification of plastid peptide transporters in silico

To identify putative plastid-localized peptide transporters *in silico* the *Arabidopsis thaliana* UniProt protein database (https://www.uniprot.org/) was interrogated with BLASTP (https://blast.ncbi.nlm.nih.gov/Blast.cgi) (Bateman et al., 2025). Predicted subcellular localization were retrieved from SUBA5 consensus dataset (Hooper et al., 2014; Hooper et al., 2017). Protein domain annotations have been performed using ScanProsite (https://prosite.expasy.org/scanprosite/), CCTOP (https://cctop.ttk.hu/) and TargetP (https://services.healthtech.dtu.dk/services/TargetP-2.0/) online tools (de Castro et al., 2006; Dobson et al., 2015; Almagro Armenteros et al., 2019). Multiple sequence alignment was performed using MUSCLE (https://www.ebi.ac.uk/Tools/msa/muscle/), phylogenetic analysis was performed using PhyML (https://toolkit.tuebingen.mpg.de/tools/phyml) and iTOL (https://itol.embl.de/) (Edgar, 2004; Guindon et al., 2010; Gabler et al., 2020; Letunic and Bork, 2024).

### Plant material and growth conditions

The Arabidopsis T-DNA insertional mutant lines described in this work *tap1-1* (SALK_085664), *nap8-1* (SALK_151551) and *ath12-1* (SALK_052673) were purchased from the Nottingham Arabidopsis Stock Centre (NASC; https://arabidopsis.info/). T-DNA insertion sites were verified by Sanger sequencing of the region of interest amplified through PCR. Higher order mutants were obtained by manual crossing and PCR-based segregation analyses of F2 populations. *ftsh2-2* (SAIL_253_A03), *ftsh11* (SALK_033047), *deg1* (Gabi-Kat_414D07) and *deg1 deg5 deg8* (cross from Gabi-Kat_414D07, SALK_099162 and SALK_004770) mutant lines were previously described (Butenko et al., 2018b; Adam et al., 2019; Tadini et al., 2020). *TAP1-*, *NAP8-* and *ATH12-GFP/-RFP* constructs were obtained by Gateway cloning (Invitrogen) using pDONR207 or pDONR201 as donor plasmids and pB2GW7 (GFP) or pB7RWG2 (RFP) as expression vectors (gatewayvectors.vib.be). Coding sequences were cloned in frame with *GFP* or *RFP* sequences at the C-termini under the control of CaMV35S constitutive promoter. Lines were obtained by Agrobacterium-mediated transformation. *TAP1-GFP TIC20-RFP*, *TAP1-GFP NAP8-RFP*, *TAP1-GFP ATH12-RFP* lines were obtained by manual crossing and segregation analysis. *Arabidopsis thaliana* Col-0 ecotype, used as control, or mutant plants were cultivated either on soil or Petri dishes in growth chambers under long day conditions (16 h at 100 μmol photons m^-2^ s^-1^ light and 8 h dark, at 22°C temperature). For heat sensitivity wild-type plants and the triple mutant ppts were surface-sterilized and sown on Murashige and Skoog (MS) medium (Duchefa, Haarlem, The Netherlands) supplemented with 2% (w/v) sucrose and 1.5% (w/v) Phyto-Agar (Duchefa). Plants were grown for 15 days at 22 °C under a 16 h light/8 h dark photoperiod and then subjected to heat stress at 45 °C for 2 h (HS). After treatment, seedlings were returned to 22 °C for either 3 h (R) or 48 h (2d-RHS) to allow physiological recovery. For morphological analyses, seeds were sown in a single row at the top of MS agar plates, which were placed vertically and maintained under the same growth conditions described above. Collected samples were immediately frozen in liquid nitrogen and stored at −80 °C until further analysis. Five biological replicates were analysed for each time point, and each experiment was independently repeated at least three times.

### Accession numbers

Arabidopsis Genome Initiative accession numbers (TAIR; https://www.arabidopsis.org) of the *A. thaliana* genes mentioned in this work: *TAP1* (AT1G70610), *NAP8* (AT4G25450), *ATH12* (AT5G03910), *FTSH2* (AT2G30950), *FTSH11* (AT5G53170), *DEG1* (AT3G27925), *DEG5* (AT4G18370), *DEG8* (AT5G39830), *TIC20-II* (AT2G47840), *CLPB3* (AT5G15450), *LHCB4* (AT5G01530), *PSBO1* (AT5G66570), *RbcL* (ATCG00490), *PSBQ2* (AT4G05180), *ABCB1* (AT2G36910), *ABCB4* (AT2G47000), *ABCB19* (AT3G28860), *APX1* (AT1G07890), *APX2* (AT3G09640), *tAPX* (AT1G77490), *FSD1* (AT4G25100), *CAT2* (AT4G35090), *CuZnSD1* (AT1G08830), *FSD2* (AT5G51100), *FSD3* (AT5G23310), *CuZnSD2* (AT2G28190), *UBQ10* (AT4G05320), *ACT8* (AT1G49240), *PP2AA3* (AT1G13320).

### Fluorescence proteins localization

Protoplast isolation was performed as previously described (Yoo et al., 2007; Costa et al., 2012). Well-expanded rosette leaves from 18 DAS were cut into strips of 1-5 mm with a scalpel blade. Leaf tissue was incubated in a buffer containing 1.25% (w/v) cellulase Onozuka R-10 (Duchefa) and 0.3% (w/v) Macerozyme R-10 (Duchefa) for 3h at room temperature in the dark. The suspension was filtered through a 50 μm nylon mesh and washed with a solution containing 154 mM NaCl, 125 mM CaCl_2_, 5 mM KCl, 2 mM MES pH 5.7. Fluorescence proteins localization was performed through an upright Nikon A1 confocal microscope using the following settings: GFP channel excitation 485 nm and detection 500-550 nm; RFP channel excitation 560 nm and detection 570-616 nm; Chlorophyll channel excitation 485 nm and detection 663-738nm.

### Complementation in yeast

*Saccharomyces cerevisiae* BY4741 (genetic background S288C) strain, kindly donated by Prof. Federico Lazzaro, was employed. For the generation of *mdl1Δ TAP1*, *mdl1Δ NAP8* and *mdl1Δ ATH12* strains, the DNA recombination cassettes were produced by Duolix cloned into pSEVA18 plasmids. Each cassette contains the coding sequence of either of *TAP1*, *NAP8* or *ATH12* with the respective chloroplast transit peptides portion substituted in frame with the *MDL1* native mitochondrial transit peptide and, downstream, the *KANMX* for selection. Each recombination cassette is flanked by 40 bp homologous to drive the replacement of the entire *MDL1*, allowing for the expression under the native promoter. *EcoRI* restriction sites were added to linearize the cassettes prior to transformation. The CDS of *NAP8* naturally bears an *EcoRI* restriction site, thus, it was edited by substituting the thymine 1416 with a cytosine to disrupt the site without altering the aminoacidic sequence. The recombination cassette employed for the generation of the *mdl1Δ* null mutant strain was obtained by High-Fidelity PCR amplification of the *KANMX* locus using the pFA6a plasmid as a template. The used primers were designed to add 40 bp homologous able to replace completely the *MDL1* locus through homologous recombination. Yeast transformation was performed as previously described (Gietz and Woods, 2002). Correct recombination was confirmed through PCR amplification and Sanger sequencing. Heat sensitivity of yeast strains was assessed as previously described with a few modifications (Jarolim et al., 2013). Yeast was grown in liquid complete medium at 28°C in shaking until the exponential phase was reached (0.5-0.6 OD600). Then, culture was sampled, serial diluted and plated on complete medium as T0 sample. After sampling, culture was halved into subcultures. One subculture was incubated at 45°C for 2h whereas the other half was kept at 28°C as control. Then, both subcultures were sampled, serial diluted and plated on complete medium as Heat and Mock samples. Petri’s dishes were incubated at 28°C for 3d to allow colonies formation. Obtained colonies were then counted to estimate the number of viable cells in Heat and Mock samples with respect to T0 sample for each strain.

### Split-Ubiquitin assay

Protein-protein interactions among TAP1, NAP8 and ATH12 in all combinations was performed using the mating-based split-ubiquitin system and procedure explained in previous works (Obrdlik et al., 2004; Grefen et al., 2007). Vectors and strains were purchased from the Arabidopsis Biological Resource Center (ABCR, https://abrc.osu.edu/). The coding sequence of either *TAP1*, *NAP8* or *ATH12* with the respective chloroplast transit peptide portion removed were cloned in frame at the C-terminus with the C- (Cub) or N-terminal (NubG) portion of ubiquitin within pMetYC-DEST (for bait-Cub fusions) and pXN21-DEST (for prey-NubG fusions) vectors, respectively, by Gateway cloning (Invitrogen). Obtained bait-Cub and prey-NubG vectors were transformed into THY.AP4 and THY.AP5 and mated to yield diploid strains suitable for the assay. Diploid strains were serial diluted and spotted on solid synthetic dextrose minimal media either lacking or supplemented with histidine and adenine for control and restrictive conditions, respectively, and incubated at 28°C for 3d to allow growth.

### Protein samples preparation and immunoblot analyses

For fresh or frozen leaf material, samples were homogenized in Laemmli buffer containing 20% glycerol, 4% (w/v) SDS, 160 mm Tris-HCl pH 6.8, 10% (v/v) 2-mercaptoethanol with a ratio of 10 µL of buffer per mg of plant material (Tadini et al., 2023). Samples were incubated at 65°C for 15 min and centrifuged for 10 min at 16000 g to pellet debris. Cleared homogenates were incubated for 5 min at 95°C. Samples were fractionated by SDS-PAGE (Schägger and von Jagow, 1987) and then blotted onto PVDF membranes. Primary antibodies raised against RbcL, CLPB3, LHCB4 and PSBO were obtained from Agrisera. Primary antibody raised against GFP was purchased from Invitrogen. Immunodetection was performed using BioRad ChemiDoc imaging system and BioRad ImageLab software.

### Chloroplast enriched fraction and sub-fractionation

Chloroplast enriched fractions were obtained as described previously (Kunst, 1998), with minor changes. 350 mg of leaves were homogenized in 4 mL of a buffer containing 45 mM sorbitol, 20 mM Tricine-KOH pH 8.4, 10 mM EDTA, 10 mM NaHCO3 and 0.1% (w/v) BSA fraction V, supplemented with cOmplete™ proteinase inhibitor cocktail (Roche), filtered through a single-layer Miracloth (Millipore) and centrifuged for 7 min at 700 g at 4°C. Pelleted chloroplasts were gently resuspended in 200 µL of extraction buffer composed of 9 mM HEPES-KOH pH 8, 0.6 mM KOAc, 0.1 mM MgOAc supplemented with cOmplete™ proteinase inhibitor cocktail (Roche). Resuspended chloroplasts were mechanically homogenized using a syringe. These preparations were aliquoted as chloroplast fractions. Subsequently, to sub-fractionate the preparations, samples were centrifugated for 10 min at 15000 g at 4 °C to pellet the chloroplast membranes. Supernatants were collected as stromal fractions, while pellets were resuspended in an equal volume of extraction buffer to obtain the membranous fractions. Samples were finally mix with an equal volume of a Laemmli buffer modified with 4% (w/v) LDS and 100 mM DTT and heated at 70°C for 20 min. Chloroplast and membrane fractions were further centrifugated for 10 min at 16000 g. Samples treated, as described above, were immediately run in SDS-PAGEs, blotted and immunodecorated with the indicated antibodies.

### Intact chloroplasts isolation and heat treatment

Intact chloroplasts were isolated from 5 g of leaves harvested from mature plants according to previous work (Kunst, 1998), with a few changes. Samples were homogenized in 250 mL of a buffer containing 45 mM sorbitol, 20 mM Tricine-KOH pH 8.4, 10 mM EDTA, 10 mM NaHCO3 and 0.1% (w/v) BSA fraction V, supplemented with cOmplete™ proteinase inhibitor cocktail (Roche), filtered through a single-layer Miracloth (Millipore) and centrifuged for 7 min at 700 g at 4°C. The pellet was gently resuspended in resuspension buffer composed of 300 mM sorbitol, 200 mM Tricine-KOH pH 8,4, 2,5 mM EDTA and 5 mM MgCl_2_. The suspension was centrifuged using a two-step percoll gradient 40%-80% (v/v) prepared with resuspension buffer at 4°C and 6500 g for 20 min in a swinging-bucket rotor (JS-13.1, Beckman Coulter). Intact chloroplasts were collected at the interface of the percoll gradient and washed once with the resuspension buffer. Samples were then normalized based on total chlorophyll amount. Samples were heat-treated at 45°C or room temperature for the indicated amount of time with a pre-heated thermoblock. Where indicated, 3 mM ATP was added to the samples just before treatment. After stress delivery, samples were centrifuged at 1000 g for 8 min. Supernatants were used for peptide quantification and identification. Pellets were used for protein degradation tests via immunoblots. Both fractions were immediately frozen in liquid nitrogen and stored at -80°C until use.

### Verification of chloroplasts integrity

Chloroplast integrity was assessed under control and heat-stress conditions via chlorophyll autofluorescence imaging. Samples were visualized using a Leica SP5 confocal microscope equipped with a 63× objective. In accordance with the protocol described by An et al. (2021), autofluorescence was detected using an excitation filter set of 510–580 nm and an emission range of 600–700 nm (An et al., 2021). Moreover, to evaluate the extent of chloroplast rupture, the relative abundance of Ribulose-1,5-bisphosphate carboxylase/oxygenase (Rubisco) was monitored in the chloroplast media before and after the heat stress. Intact chloroplasts were removed from the medium via centrifugation (3000 g for 5 min at 4°C). The resulting supernatants, containing proteins leaked from damaged organelles, were resolved by SDS-PAGE using standard manufacturer protocols (Bio-Rad). Gels were subsequently scanned with a GS800 Calibrated Densitometer. Relative quantification of the Rubisco large subunit (rbcL, ∼50 kDa) was performed using ImageJ software (NIH, USA). All experiments were carried out with at least three independent biological replicates. Statistical significance was assessed using an unpaired t-test, with a significance threshold established at p<0.05.

### Peptide purification, quantification and identification

Chloroplasts were sedimented from the media by centrifugation at 3,000×g for 5 min at 4°C. The resulting supernatants were filtered using Amicon® Ultra-0.5 Centrifugal Filter Devices (MilliporeSigma) to isolate the peptide fraction according to the manufacturer’s instructions. The samples were subsequently purified and concentrated using an SPE (Solid Phase Extraction) Clean-up Kit (Waters), in accordance with the manufacturer’s protocols. The eluted peptide fractions were evaporated to dryness under a gentle stream of nitrogen gas and subsequently reconstituted in Milli-Q® water (MilliporeSigma) for further analysis. The purified peptides were quantified via UV spectrometry at 280 nm using a calibration curve (0-10 mg/ml) realised with MS Compatible Yeast and Human Protein Extracts (Promega) as a reference. All experiments were carried out with at least three independent biological replicates. Statistical significance was assessed using an unpaired t-test, with a significance threshold established at p<0.05.

Peptide identification was performed using a a QExactive mass spectrometer coupled to a nano EasyLC 1000 (Thermo Fisher Scientific Inc.). The mobile phases consisted of 0.1% formic acid (A) and 0.1% formic acid, 99.9% acetonitrile (B). Samples (4 μL of peptides) were injected onto a self-packed ReproSil-Pur 120 C18-AQ (1.9 μm) column (75 μm × 150 mm; Dr. Maisch GmbH). A flow rate of 300 nL/min was maintained with a gradient of 2–35% B over 80 min, followed by 47% B in 4 min, and 98% B in 4 min. Sample acquisition was randomized. The mass spectrometer operated in data-dependent acquisition (DDA) mode. Full-scan MS spectra were acquired from 300–1700 m/z, and the twelve most intense signals per cycle were fragmented. HCD spectra were obtained at a resolution of 35000 using a normalized collision energy of 25 and a maximum injection time of 120 ms. The automatic gain control (AGC) target was 50,000 ions. Charge state screening was enabled, rejecting singly and unassigned charge states. Precursor masses previously selected were excluded for 30 s with a 10 ppm exclusion window. Internal lock mass calibration was applied using m/z 371.1010 and 445.1200. Three independent biological replicas were performed. Raw mass spectrometry data were then processed using MaxQuant software (version 2.6.7.0). To enhance protein identification, the "match between runs" (MBR) feature was enabled with a matching time window of 0.7 min. Tandem MS spectra were searched against a combined *Arabidopsis thaliana* protein database (UniProt/TAIR11, accessed May 2025). Given the nature of the samples, an unspecific digestion mode was employed. Variable modifications included methionine oxidation and N-terminal acetylation. Initial search tolerances were set to 20 ppm for peptides and 0.5 Da for MS/MS fragments. A 1% false discovery rate (FDR) was strictly applied at the peptide-spectrum match (PSM) level, and only peptides with a minimum length of five amino acids were retained. Comparative analysis of unique and shared peptides between experimental conditions was performed and visualized using Venn diagrams.

The physicochemical properties and potential antibacterial activity of the identified peptides were evaluated using the Database of Antimicrobial Activity and Structure of Peptides (DBAASP) v3.0 (Pirtskhalava et al., 2021). General antibacterial potential was predicted via machine learning algorithms based on the peptides’ intrinsic physicochemical characteristics (Vishnepolsky and Pirtskhalava, 2014). Peptide antioxidant properties were assessed using the Antioxidative Peptide Predictor (AnOxPP-V1.0) (Qin et al., 2023). Subcellular localisation of source proteins was determined interrogating SUBA5 and The Plant Proteome Database (http://ppdb.tc.cornell.edu/) (Sun et al., 2009).

### Protein degradation rate detection

Leaf samples were harvested from mature plants and vacuum infiltrated (3x cycles of 45 s each) in 1 ml infiltration buffer composed of 0.4% (w/v) MS salt, 0.05% Tween20 and 25 µg/mL cycloheximide and then incubated at room temperature for 30 min as pre-treatment. Next, samples were incubated at 45 °C, or room temperature, for 2h. After treatment, buffer was removed and the samples were immediately frozen in liquid nitrogen and stored at -80 °C until processing for immunoblot analysis.

### Photosynthetic pigments quantification

For pigment quantification, leaf samples (50 mg) were ground in liquid nitrogen with 80% acetone (1:20 w/v) and the homogenates centrifuged at 20,000 g for 20 minutes at 4°C. The supernatant absorbances at 663.2, 646.8, and 470 nm were measured spectrophotometrically as previously performed (ZHANG and KIRKHAM, 1996). Concentration of chlorophyll a (Chl a) and chlorophyll b (Chl b), as well as total carotenoids, expressed as mg g^-1^ fresh weight, were calculated according to previous work (Lichtenthaler, 1987):

Chl A =12,25A*663.2 - 2,79*A646.8
Chl B =21,50*A646.8 - 5,10*A663.2
Carotenoids= (1000*A470-1,82 Chl A- 85,02*Chl B)/198

### Nucleic acid analyses

otal RNA was extracted using the Nippon Genetics Kit according to the manufacturer’s protocol, starting from 50 mg (fresh weight) of leaf material. RNA concentration was determined by measuring absorbance at 260 nm using a NanoDrop 2000 spectrophotometer (Thermo Fisher Scientific, USA). Complementary DNA (cDNA) was synthesized from 2 µg of total RNA using the iScript™ cDNA Synthesis Kit (Bio-Rad), following the manufacturer’s instructions. Gene expression analysis by quantitative real-time PCR (qRT-PCR) was performed using the CFX Connect Real-Time PCR Detection System (Bio-Rad, Hercules, CA, USA), employing 37.5 ng of cDNA per reaction and SsoAdvanced Universal SYBR Green Supermix (Bio-Rad), according to the manufacturer’s guidelines. *UBQ10* (AT4G05320), *ACT8* (AT1G49240) and *PP2AA3* (AT1G13320) were used as housekeeping genes. Primers for quantitative real-time PCR (qRT-PCR) were designed as described in Lasorella et al. (2022). Primer sequences used for quantitative PCR (qPCR) analyses are reported in Table S8. Relative quantification was performed according to the comparative Ct (threshold cycle) method (2^−DDCt^) (Livak and Schmittgen, 2001).

### Determination of oxidative markers and analysis of enzymatic and non-enzymatic antioxidants

The level of lipid peroxidation was assessed by measuring malondialdehyde (MDA) content using the thiobarbituric acid (TBA) reaction, as described (Paradiso et al., 2008). The concentration of the MDA–TBA complex was calculated using an extinction coefficient of 155 mM⁻¹ cm⁻¹. Protein oxidation was evaluated spectrophotometrically by measuring the carbonyl group content following reaction with dinitrophenylhydrazine (DNPH), as reported (Romero-Puertas et al., 2002). Carbonyl content was calculated using an extinction coefficient of 22 mM⁻¹ cm⁻¹. For ascorbate (ASC) and glutathione (GSH) analysis, 0.3 g of seedlings were homogenized at 4°C with 5% methaphosphoric acid (1:4 w/v). After centrifugation at 18,000 x g for 20 minutes, the supernatants were collected, and ASC and GSH levels were determined using the colorimetric assay described (de Pinto et al., 1999). The in vitro antioxidant activity of the three peptides was evaluated by measuring their DPPH and ABTS radical scavenging activities, as described (Zhao and Liu, 2023). In both assays, the antioxidant activity was expressed as a percentage of radical scavenging activity. Peptides sequences were determined among those detected exclusively in wild type heat-stress condition, generated by the highly represented proteins PSBO1 and LHCB4 (Table S5). Peptides were synthesised by Bio-Fab Research s.r.l. (https://www.biofabresearch.com/it/). For quantifying the enzymatic antioxidant activities, samples were ground to a fine powder in liquid nitrogen and mixed with extraction buffer (1:4 w/v) containing 50 mM Tris-HCl, pH 7.5, 0.05% (w/v) cysteine, 0.1% bovine serum albumin, and 1 mM phenylmethanesulfonyl fluoride. To determine ascorbate peroxidase activity, 1 mM ASC was added to the buffer. After centrifugation at 20,000 x g for 20 minutes at 4°C, the supernatants were used for the spectrophotometric analysis. Superoxide dismutase (SOD, EC 1.15.1.1), catalase (CAT, EC 1.11.1.6), and Ascorbate peroxidase (APX, EC 1.11.1.11) activities were spectrophotometrically determined according to the methods described (Paradiso et al., 2020).

## Author contributions

NJ, GD, LT, CV and PP conceptualized the study. NJ performed the in silico identification of plastid peptide transporters. NJ and LT took care of Arabidopsis lines generation and isolation. NJ and LT performed subcellular localization experiments. CB performed Split-Ub analysis. NJ performed yeast complementation analyses. NJ, GD, EC and LT performed chloroplasts isolation and treatments. GD and EC performed peptide quantification and MS analysis. NJ, LT and MR performed immunoblot analyses. MCdP, SF, CL and SV performed phenotypic analysis on heat-treated plant material and took care of qRT-PCR and the determination of oxidative markers and analysis of enzymatic and non-enzymatic antioxidants. NJ, GD, LT, CV, SF, MCdP contributed to the drafting of the manuscript. NJ, GD, LT, CV, MCdP and PP oversaw the study and finalized the manuscript. All authors approved its publication and accepted responsibility for the study.

## Acknowledgment and funding

We are grateful to Manuel Rodríguez-Conceptíon, Alessandro Vitale, Anna Moroni, Michela Zottini and Tomas Morosinotto for fruitful conversations and suggestions. We are also grateful to Valerio Paravicini and Mario Beretta for technical assistance. Part of this work was carried out at NOLIMITS, an advanced imaging facility at the Università degli Studi di Milano. This study was carried out with the support of MUR—Ministero dell’Università e della Ricerca, grant number PRIN 2017-FBS8YN, awarded to PP.

## Figure legends

**Figure S1 Characterization of TAP1, NAP8, and ATH12 mutant lines. A)** Schematic representation of the genomic organization of the TAP1, NAP8, and ATH12 loci. Exons are shown as white boxes and introns as black lines. The positions of T-DNA insertions, as well as translation start and stop codons, are indicated. Gene models are drawn to scale, whereas T-DNA insertions are not. **B)** Representative relative transcript levels of *TAP1*, *NAP8*, *NAP8 T-DNA* and *ATH12* loci. Bars represent standard errors. **C)** Representative images of 15 DAS plants of the indicated genotypes grown under long-day photoperiod conditions in soil under controlled growth chamber settings.

**Figure S2 Supplementary data on peptide extrusion**. **A**) Representative confocal images of purified wild-type and *tap1 nap8 ath12* chloroplasts before and after 60 min incubation at 45°C. **B**) Representative Coomassie Brilliant Blue–stained SDS–PAGE gel showing the Rubisco large subunit (RbcL) detected in supernatants from Col-0 and *tap1 nap8 ath12* chloroplasts under control conditions and after 30 and 60 min of heat treatment. **C**) Frequency distribution of peptide lengths (in amino acids) recovered from the incubation buffer of chloroplasts treated at 45°C in the presence of ATP, based on LC–MS/MS analyses from at least three independent experiments. Data are normalized to the total MS signal intensity of the corresponding genotype. **D**) Relative abundance of peptides detected in the incubation buffer of wild-type chloroplasts treated at 45°C in the presence of ATP, grouped according to the plastid or non-plastid localization of their source proteins. Percentages refer to plastid-derived proteins. Data are derived from three independent LC–MS/MS experiments. **E)** Distribution of unique Col-0 heat-induced peptides according to their net charge. Values were normalized to MS signal intensity. **F)** Distribution of unique Col-0 heat-induced peptides based on their Moon and Fleming hydrophobicity index. Values were normalized to MS signal intensity. **G**) Sequence logos generated using Seq2Logo 2.0 from peptides identified in Col-0 heat-treated supernatants, illustrating amino acid preferences within the N-terminal (upper panel) and C-terminal (lower panel) cleavage windows. The height of each letter represents its relative frequency or information content at each position. Red dashed lines indicate cleavage sites. For the N-terminal logo, P1 corresponds to the residue preceding the cleavage site and P–1 to the residue following it; for the C-terminal logo, P1′ indicates the residue preceding the cleavage site and P–1′ the residue following it.

**Figure S3 Multiple sequence alignment of major source proteins generating Col-0 heat-induced peptides. A)** Multiple sequence alignment of Col-0 heat-induced peptide sequences mapped onto the PSBO1 (A-B), PSBQ2 (C-D) and LHCB4 (E) protein sequences, delineating the peptide-generating region.

**Figure S4 Expression analysis of major antioxidant enzymes. A–I)** Relative transcript levels of the indicated genes measured in plantlets before heat treatment (Control), immediately after heat treatment (45°C for 2 h), and after 3 h of recovery. Different letters indicate statistically significant differences according to one-way ANOVA followed by Tukey–Kramer post hoc test (*P* < 0.05). Error bars represent the standard errors of at least three independent biological replicates.

**Table S1 BLASTP search results for candidate peptide transporters.** Lists of significant BLASTP hits obtained using the mitochondrial peptide exporters HAF-1 from *Caenorhabditis elegans* and Mdl1 from *Saccharomyces cerevisiae* as query sequences against the *Arabidopsis thaliana* reference protein database.

**Table S2 Identified peptide sequences in chloroplast supernatants.** List of peptide sequences identified by LC–MS/MS under each experimental condition from three independent biological replicates.

**Table S3 Source proteins of peptides identified in chloroplast supernatants.** List of proteins from which the identified peptide sequences were derived, as determined by LC–MS/MS analysis.

**Table S4 Proteins giving rise to chloroplast-extruded peptides.** List of proteins identified as sources of chloroplast-extruded peptides, including Araport ID, UniProt ID, gene name, subcellular localization, protein name, peptide counts from two independent experiments, mean peptide count, and relative abundance expressed as a percentage. The Excel file contains two worksheets: the first includes all identified proteins, whereas the second is restricted to plastid-localized proteins.

**Table S5 Col-0 heat-induced peptides and their source proteins.** List of peptide sequences specifically induced by heat treatment in Col-0, reporting the Araport ID, UniProt ID, and functional description of the source proteins, together with the main chemical and physicochemical parameters of each peptide.

**Table S6 Comparison of physicochemical properties of peptide pools.** Relevant physicochemical properties of secreted peptide pools from chloroplasts (this work) and mitochondria (Young et al., 2001; Augustin et al., 2005) are listed for comparison.

**Table S7 List of peptide core sequences.** List of conserved core peptide sequences identified within peptides derived from PSBO1, PSBQ2 and LHCB4, together with their frequency of occurrence and positions within the protein sequences.

**Table S8 List of primers used in this work.** List of primer sequences employed for genotyping, qRT-PCR, cloning and yeast homologous recombination.

## References

Abele, R., & Tampé, R. (2018). Moving the Cellular Peptidome by Transporters. Frontiers in Cell and Developmental Biology, 6. 10.3389/fcell.2018.00043

Adam, Z., Aviv-Sharon, E., Keren-Paz, A., Naveh, L., Rozenberg, M., Savidor, A., & Chen, J. (2019). The chloroplast envelope protease FTSH11 – interaction with CPN60 and identification of potential substrates. Frontiers in Plant Science, 10, 428. 10.3389/fpls.2019.00428

Allakhverdiev, S. I., Kreslavski, V. D., Klimov, V. V., Los, D. A., Carpentier, R., & Mohanty, P. (2008). Heat stress: An overview of molecular responses in photosynthesis. In Photosynthesis Research (Vol. 98, Numbers 1–3, pp. 541–550). Springer. 10.1007/s11120-008-9331-0

Almagro Armenteros, J. J., Salvatore, M., Emanuelsson, O., Winther, O., von Heijne, G., Elofsson, A., & Nielsen, H. (2019). Detecting sequence signals in targeting peptides using deep learning. Life Science Alliance, 2(5), e201900429. 10.26508/lsa.201900429

An, J., Miao, X., Wang, L., Li, X., Liu, X., & Gao, H. (2021). Visualizing the Integrity of Chloroplast Envelope by Rhodamine and Nile Red Staining. Frontiers in Plant Science, 12. 10.3389/fpls.2021.668414

Apel, K., & Hirt, H. (2004). REACTIVE OXYGEN SPECIES: Metabolism, Oxidative Stress, and Signal Transduction. Annu. Rev. Plant Biol, 55, 373–399. 10.1146/annurev.arplant.55.031903.141701

Augustin, S., Nolden, M., Müller, S., Hardt, O., Arnold, I., & Langer, T. (2005). Characterization of peptides released from mitochondria. Journal of Biological Chemistry, 280(4), 2691–2699. 10.1074/jbc.M410609200

Bateman, A., Martin, M.-J., Orchard, S., Magrane, M., Adesina, A., Ahmad, S., Bowler-Barnett, E. H., Bye-A-Jee, H., Carpentier, D., Denny, P., Fan, J., Garmiri, P., Gonzales, L. J. da C., Hussein, A., Ignatchenko, A., Insana, G., Ishtiaq, R., Joshi, V., Jyothi, D., … Zhang, J. (2025). UniProt: the Universal Protein Knowledgebase in 2025. Nucleic Acids Research, 53(D1), D609–D617. 10.1093/nar/gkae1010

Butenko, Y., Lin, A., Naveh, L., Kupervaser, M., Levin, Y., Reich, Z., & Adam, Z. (2018a). Differential Roles of the Thylakoid Lumenal Deg Protease Homologs in Chloroplast Proteostasis. Plant Physiology, 178(3), 1065–1080. 10.1104/pp.18.00912

Butenko, Y., Lin, A., Naveh, L., Kupervaser, M., Levin, Y., Reich, Z., & Adam, Z. (2018b). Differential Roles of the Thylakoid Lumenal Deg Protease Homologs in Chloroplast Proteostasis. Plant Physiology, 178(3), 1065–1080. 10.1104/pp.18.00912

Chen, L., Li, Q., & Li, L. (2014). The modeled structures of Deg5 and Deg8 proteases in Arabidopsis thaliana. TURKISH JOURNAL OF BIOLOGY, 38, 168–176. 10.3906/biy-1306-74

Chloupková, M., LeBard, L. S., & Koeller, D. M. (2003). MDL1 is a High Copy Suppressor of ATM1: Evidence for a Role in Resistance to Oxidative Stress. Journal of Molecular Biology, 331(1), 155–165. 10.1016/S0022-2836(03)00666-1

Costa, A., Gutla, P. V. K., Boccaccio, A., Scholz-Starke, J., Festa, M., Basso, B., Zanardi, I., Pusch, M., Schiavo, F. Lo, Gambale, F., & Carpaneto, A. (2012). The *Arabidopsis* central vacuole as an expression system for intracellular transporters: functional characterization of the Cl ^−^ /H ^+^ exchanger CLC-7. The Journal of Physiology, 590(15), 3421–3430. 10.1113/jphysiol.2012.230227

Czechowski, T., Stitt, M., Altmann, T., Udvardi, M. K., & Scheible, W.-R. (2005). Genome-Wide Identification and Testing of Superior Reference Genes for Transcript Normalization in Arabidopsis. Plant Physiology, 139(1), 5–17. 10.1104/pp.105.063743

Davidson, A. L., Dassa, E., Orelle, C., & Chen, J. (2008). Structure, Function, and Evolution of Bacterial ATP-Binding Cassette Systems. Microbiology and Molecular Biology Reviews, 72(2), 317–364. 10.1128/MMBR.00031-07

de Castro, E., Sigrist, C. J. A., Gattiker, A., Bulliard, V., Langendijk-Genevaux, P. S., Gasteiger, E., Bairoch, A., & Hulo, N. (2006). ScanProsite: detection of PROSITE signature matches and ProRule-associated functional and structural residues in proteins. Nucleic Acids Research, 34(Web Server), W362–W365. 10.1093/nar/gkl124

De Las Rivas, J., & Barber, J. (2004). Analysis of the Structure of the PsbO Protein and its Implications. Photosynthesis Research, 81(3), 329–343. 10.1023/B:PRES.0000036889.44048.e4

Detmers, F. J. M., Lanfermeijer, F. C., & Poolman, B. (2001). Peptides and ATP binding cassette peptide transporters. Research in Microbiology, 152(3–4), 245–258. 10.1016/S0923-2508(01)01196-2

Dobson, L., Reményi, I., & Tusnády, G. E. (2015). CCTOP: a Consensus Constrained TOPology prediction web server. Nucleic Acids Research, 43(W1), W408–W412. 10.1093/nar/gkv451

Dogra, V., Li, M., Singh, S., Li, M., & Kim, C. (2019). Oxidative post-translational modification of EXECUTER1 is required for singlet oxygen sensing in plastids. Nature Communications, 10(1), 2834. 10.1038/s41467-019-10760-6

Edgar, R. C. (2004). MUSCLE: A multiple sequence alignment method with reduced time and space complexity. BMC Bioinformatics, 5(1), 1–19. 10.1186/1471-2105-5-113/FIGURES/16

Ferro, E. S., Rioli, V., Castro, L. M., & Fricker, L. D. (2014). Intracellular peptides: From discovery to function. EuPA Open Proteomics, 3, 143–151. 10.1016/j.euprot.2014.02.009

Ferro, M., Brugière, S., Salvi, D., Seigneurin-Berny, D., Court, M., Moyet, L., Ramus, C., Miras, S., Mellal, M., Le Gall, S., Kieffer-Jaquinod, S., Bruley, C., Garin, J., Joyard, J., Masselon, C., & Rolland, N. (2010). AT_CHLORO, a Comprehensive Chloroplast Proteome Database with Subplastidial Localization and Curated Information on Envelope Proteins. Molecular & Cellular Proteomics. 10.1074/mcp.M900325-MCP200

Fiorese, C. J., Schulz, A. M., Lin, Y.-F., Rosin, N., Pellegrino, M. W., & Haynes, C. M. (2016). The Transcription Factor ATF5 Mediates a Mammalian Mitochondrial UPR. Current Biology, 26(15), 2037–2043. 10.1016/j.cub.2016.06.002

Fortunato, S., Lasorella, C., Dipierro, N., Vita, F., & de Pinto, M. C. (2023). Redox Signaling in Plant Heat Stress Response. Antioxidants, 12(3), 605. 10.3390/antiox12030605

Gabler, F., Nam, S., Till, S., Mirdita, M., Steinegger, M., Söding, J., Lupas, A. N., & Alva, V. (2020). Protein Sequence Analysis Using the MPI Bioinformatics Toolkit. Current Protocols in Bioinformatics, 72(1). 10.1002/cpbi.108

Grefen, C., Lalonde, S., & Obrdlik, P. (2007). Split-Ubiquitin System for Identifying Protein-Protein Interactions in Membrane and Full-Length Proteins. Current Protocols in Neuroscience, 41(1). 10.1002/0471142301.ns0527s41

Guindon, S., Dufayard, J.-F., Lefort, V., Anisimova, M., Hordijk, W., & Gascuel, O. (2010). New Algorithms and Methods to Estimate Maximum-Likelihood Phylogenies: Assessing the Performance of PhyML 3.0. Systematic Biology, 59(3), 307–321. 10.1093/sysbio/syq010

Haynes, C. M., Yang, Y., Blais, S. P., Neubert, T. A., & Ron, D. (2010). The Matrix Peptide Exporter HAF-1 Signals a Mitochondrial UPR by Activating the Transcription Factor ZC376.7 in C. elegans. Molecular Cell, 37(4), 529–540. 10.1016/j.molcel.2010.01.015

Hendrix, S., Dard, A., Meyer, A. J., & Reichheld, J.-P. (2023). Redox-mediated responses to high temperature in plants. Journal of Experimental Botany, 74(8), 2489–2507. 10.1093/jxb/erad053

Hooper, C. M., Castleden, I. R., Tanz, S. K., Aryamanesh, N., & Millar, A. H. (2017). SUBA4: the interactive data analysis centre for Arabidopsis subcellular protein locations. Nucleic Acids Research, 45(D1), D1064–D1074. 10.1093/nar/gkw1041

Hooper, C. M., Tanz, S. K., Castleden, I. R., Vacher, M. A., Small, I. D., & Millar, A. H. (2014). SUBAcon: a consensus algorithm for unifying the subcellular localization data of the *Arabidopsis* proteome. Bioinformatics, 30(23), 3356–3364. 10.1093/bioinformatics/btu550

Huesgen, P. F., Schuhmann, H., & Adamska, I. (2009). Deg/HtrA proteases as components of a network for photosystem II quality control in chloroplasts and cyanobacteria. Research in Microbiology, 160(9), 726–732. 10.1016/j.resmic.2009.08.005

Hwang, J.-U., Song, W.-Y., Hong, D., Ko, D., Yamaoka, Y., Jang, S., Yim, S., Lee, E., Khare, D., Kim, K., Palmgren, M., Yoon, H. S., Martinoia, E., & Lee, Y. (2016). Plant ABC Transporters Enable Many Unique Aspects of a Terrestrial Plant’s Lifestyle. Molecular Plant, 9(3), 338–355. 10.1016/j.molp.2016.02.003

Inaba, T., & Schnell, D. J. (2008). Protein trafficking to plastids: one theme, many variations. Biochemical Journal, 413(1), 15–28. 10.1042/BJ20080490

Jarolim, S., Ayer, A., Pillay, B., Gee, A. C., Phrakaysone, A., Perrone, G. G., Breitenbach, M., & Dawes, I. W. (2013). *Saccharomyces cerevisiae* Genes Involved in Survival of Heat Shock. G3 Genes|Genomes|Genetics, 3(12), 2321–2333. 10.1534/g3.113.007971

Jeran, N., Mercier, M., Pesaresi, P., & Tadini, L. (2025). Proteostasis and protein quality control in chloroplasts: mechanisms and novel insights related to protein mislocalization. Journal of Experimental Botany. 10.1093/jxb/eraf182

Kang, J., Park, J., Choi, H., Burla, B., Kretzschmar, T., Lee, Y., & Martinoia, E. (2011). Plant ABC Transporters. The Arabidopsis Book, 9, e0153. 10.1199/tab.0153

Kapri-Pardes, E., Naveh, L., & Adam, Z. (2007). The thylakoid lumen protease Deg1 is involved in the repair of Photosystem II from photoinhibition in *Arabidopsis*. The Plant Cell, 19(3), 1039–1047. 10.1105/tpc.106.046573

Kato, Y., & Sakamoto, W. (2018). FtsH Protease in the Thylakoid Membrane: Physiological Functions and the Regulation of Protease Activity. Frontiers in Plant Science, 9. 10.3389/fpls.2018.00855

Khavinson, V., Diomede, F., Mironova, E., Linkova, N., Trofimova, S., Trubiani, O., Caputi, S., & Sinjari, B. (2020). AEDG Peptide (Epitalon) Stimulates Gene Expression and Protein Synthesis during Neurogenesis: Possible Epigenetic Mechanism. Molecules, 25(3), 609. 10.3390/molecules25030609

Khavinson, V. K., Popovich, I. G., Linkova, N. S., Mironova, E. S., & Ilina, A. R. (2021). Peptide Regulation of Gene Expression: A Systematic Review. Molecules, 26(22), 7053. 10.3390/molecules26227053

Kreis, E., Niemeyer, J., Merz, M., Scheuring, D., & Schroda, M. (2023). CLPB3 is required for the removal of chloroplast protein aggregates and thermotolerance in *Chlamydomonas*. Journal of Experimental Botany, 74(12), 3714–3728. 10.1093/jxb/erad109

Krojer, T., Pangerl, K., Kurt, J., Sawa, J., Stingl, C., Mechtler, K., Huber, R., Ehrmann, M., & Clausen, T. (2008). Interplay of PDZ and protease domain of DegP ensures efficient elimination of misfolded proteins. Proceedings of the National Academy of Sciences, 105(22), 7702–7707. 10.1073/pnas.0803392105

Lebeau, J., Rainbolt, T. K., & Wiseman, R. L. (2018). Coordinating Mitochondrial Biology Through the Stress-Responsive Regulation of Mitochondrial Proteases. In International Review of Cell and Molecular Biology (Vol. 340, pp. 79–128). 10.1016/bs.ircmb.2018.05.003

Lehnert, E., Mao, J., Mehdipour, A. R., Hummer, G., Abele, R., Glaubitz, C., & Tampé, R. (2016). Antigenic Peptide Recognition on the Human ABC Transporter TAP Resolved by DNP-Enhanced Solid-State NMR Spectroscopy. Journal of the American Chemical Society, 138(42), 13967–13974. 10.1021/jacs.6b07426

Letunic, I., & Bork, P. (2024). Interactive Tree of Life (iTOL) v6: recent updates to the phylogenetic tree display and annotation tool. Nucleic Acids Research, 52(W1), W78–W82. 10.1093/nar/gkae268

Lewis, V. G., Ween, M. P., & McDevitt, C. A. (2012). The role of ATP-binding cassette transporters in bacterial pathogenicity. Protoplasma, 249(4), 919–942. 10.1007/s00709-011-0360-8

Liesa, M., Qiu, W., & Shirihai, O. S. (2012). Mitochondrial ABC transporters function: The role of ABCB10 (ABC-me) as a novel player in cellular handling of reactive oxygen species. Biochimica et Biophysica Acta - Molecular Cell Research, 1823(10), 1945–1957. 10.1016/j.bbamcr.2012.07.013

Lin, R., & Wang, H. (2005). Two Homologous ATP-Binding Cassette Transporter Proteins, AtMDR1 and AtPGP1, Regulate Arabidopsis Photomorphogenesis and Root Development by Mediating Polar Auxin Transport. Plant Physiology, 138(2), 949–964. 10.1104/PP.105.061572

Liu, K., Zhao, H., Lee, K. P., Yu, Q., Di, M., Wang, L., & Kim, C. (2024). EXECUTER1 and singlet oxygen signaling: A reassessment of nuclear activity. The Plant Cell, 37(1). 10.1093/plcell/koae296

Locher, K. P. (2016). Mechanistic diversity in ATP-binding cassette (ABC) transporters. Nature Structural & Molecular Biology, 23(6), 487–493. 10.1038/nsmb.3216

Luciński, R., & Adamiec, M. (2023). The role of plant proteases in the response of plants to abiotic stress factors. Frontiers in Plant Physiology, 1. 10.3389/fphgy.2023.1330216

Lyapina, I., Ivanov, V., & Fesenko, I. (2021). Peptidome: Chaos or Inevitability. International Journal of Molecular Sciences, 22(23), 13128. 10.3390/ijms222313128

Monnet, V. (2003). Bacterial oligopeptide-binding proteins. Cellular and Molecular Life Sciences (CMLS), 60(10), 2100–2114. 10.1007/s00018-003-3054-3

Myouga, F., Motohashi, R., Kuromori, T., Nagata, N., & Shinozaki, K. (2006). An Arabidopsis chloroplast-targeted Hsp101 homologue, APG6, has an essential role in chloroplast development as well as heat-stress response. Plant Journal, 48(2), 249–260. 10.1111/j.1365-313X.2006.02873.x

Nijenhuis, M., & Hämmerling, G. J. (1996). Multiple regions of the transporter associated with antigen processing (TAP) contribute to its peptide binding site. Journal of Immunology (Baltimore, Md. : 1950), 157(12), 5467–5477. http://www.ncbi.nlm.nih.gov/pubmed/8955196

Nishimura, K., Kato, Y., & Sakamoto, W. (2017). Essentials of Proteolytic Machineries in Chloroplasts. Molecular Plant, 10(1), 4–19. 10.1016/j.molp.2016.08.005

Noh, B., Murphy, A. S., & Spalding, E. P. (2001). Multidrug Resistance-Like Genes of Arabidopsis Required for Auxin Transport and Auxin-Mediated Development. The Plant Cell, 13(11), 2441. 10.2307/3871586

Nöll, A., Thomas, C., Herbring, V., Zollmann, T., Barth, K., Mehdipour, A. R., Tomasiak, T. M., Brüchert, S., Joseph, B., Abele, R., Oliéric, V., Wang, M., Diederichs, K., Hummer, G., Stroud, R. M., Pos, K. M., & Tampé, R. (2017). Crystal structure and mechanistic basis of a functional homolog of the antigen transporter TAP. Proceedings of the National Academy of Sciences, 114(4), E438–E447. 10.1073/pnas.1620009114

Obrdlik, P., El-Bakkoury, M., Hamacher, T., Cappellaro, C., Vilarino, C., Fleischer, C., Ellerbrok, H., Kamuzinzi, R., Ledent, V., Blaudez, D., Sanders, D., Revuelta, J. L., Boles, E., André, B., & Frommer, W. B. (2004). K ^+^ channel interactions detected by a genetic system optimized for systematic studies of membrane protein interactions. Proceedings of the National Academy of Sciences, 101(33), 12242–12247. 10.1073/pnas.0404467101

Parcerisa, I. L., Rosano, G. L., & Ceccarelli, E. A. (2020). Biochemical characterization of ClpB3, a chloroplastic disaggregase from Arabidopsis thaliana. Plant Molecular Biology, 104(4–5), 451–465. 10.1007/s11103-020-01050-7

Paul, P., Mesihovic, A., Chaturvedi, P., Ghatak, A., Weckwerth, W., Böhmer, M., & Schleiff, E. (2020). Structural and Functional Heat Stress Responses of Chloroplasts of Arabidopsis thaliana. Genes, 11(6), 650. 10.3390/genes11060650

Pirtskhalava, M., Amstrong, A. A., Grigolava, M., Chubinidze, M., Alimbarashvili, E., Vishnepolsky, B., Gabrielian, A., Rosenthal, A., Hurt, D. E., & Tartakovsky, M. (2021). DBAASP v3: database of antimicrobial/cytotoxic activity and structure of peptides as a resource for development of new therapeutics. Nucleic Acids Research, 49(D1), D288–D297. 10.1093/nar/gkaa991

Qin, D., Jiao, L., Wang, R., Zhao, Y., Hao, Y., & Liang, G. (2023). Prediction of antioxidant peptides using a quantitative structure−activity relationship predictor (AnOxPP) based on bidirectional long short-term memory neural network and interpretable amino acid descriptors. Computers in Biology and Medicine, 154, 106591. 10.1016/j.compbiomed.2023.106591

Quirós, P. M., Mottis, A., & Auwerx, J. (2016). Mitonuclear communication in homeostasis and stress. Nature Reviews Molecular Cell Biology, 17(4), 213–226. 10.1038/nrm.2016.23

Rees, D. C., Johnson, E., & Lewinson, O. (2009). ABC transporters: the power to change. Nature Reviews Molecular Cell Biology, 10(3), 218–227. 10.1038/nrm2646

Roose, J. L., Kashino, Y., & Pakrasi, H. B. (2007). The PsbQ protein defines cyanobacterial Photosystem II complexes with highest activity and stability. Proceedings of the National Academy of Sciences, 104(7), 2548–2553. 10.1073/pnas.0609337104

Sánchez-Fernández, R., Davies, T. G. E., Coleman, J. O. D., & Rea, P. A. (2001). The Arabidopsis thaliana ABC Protein Superfamily, a Complete Inventory. Journal of Biological Chemistry, 276(32), 30231–30244. 10.1074/jbc.M103104200

Santelia, D., Vincenzetti, V., Azzarello, E., Bovet, L., Fukao, Y., Düchtig, P., Mancuso, S., Martinoia, E., & Geisler, M. (2005). MDR-like ABC transporter AtPGP4 is involved in auxin-mediated lateral root and root hair development. FEBS Letters, 579(24), 5399–5406. 10.1016/J.FEBSLET.2005.08.061

Schaedler, T. A., Faust, B., Shintre, C. A., Carpenter, E. P., Srinivasan, V., van Veen, H. W., & Balk, J. (2015). Structures and functions of mitochondrial ABC transporters. Biochemical Society Transactions, 43(5), 943–951. 10.1042/BST20150118

Schägger, H., & von Jagow, G. (1987). Tricine-sodium dodecyl sulfate-polyacrylamide gel electrophoresis for the separation of proteins in the range from 1 to 100 kDa. Analytical Biochemistry, 166(2), 368–379. 10.1016/0003-2697(87)90587-2

Schuhmann, H., Huesgen, P. F., Gietl, C., & Adamska, I. (2008). The DEG15 Serine Protease Cleaves Peroxisomal Targeting Signal 2-Containing Proteins in Arabidopsis. Plant Physiology, 148(4), 1847–1856. 10.1104/pp.108.125377

Staacke, T., Mueller-Roeber, B., & Balazadeh, S. (2025). Stress resilience in plants: the complex interplay between heat stress memory and resetting. New Phytologist, 245(6), 2402–2421. 10.1111/nph.20377

Stieger, B., & Higgins, C. F. (2007). Twenty years of ATP-binding cassette (ABC) transporters. Pflügers Archiv - European Journal of Physiology, 453(5), 543–543. 10.1007/s00424-006-0159-1

Sun, Q., Zybailov, B., Majeran, W., Friso, G., Olinares, P. D. B., & van Wijk, K. J. (2009). PPDB, the Plant Proteomics Database at Cornell. Nucleic Acids Research, 37(suppl_1), D969–D974. 10.1093/nar/gkn654

Sun, X., Ouyang, M., Guo, J., Ma, J., Lu, C., Adam, Z., & Zhang, L. (2010). The thylakoid protease Deg1 is involved in photosystem-II assembly in Arabidopsis thaliana. The Plant Journal, 62(2), 240–249. 10.1111/j.1365-313X.2010.04140.x

Tadini, L., Jeran, N., Domingo, G., Zambelli, F., Masiero, S., Calabritto, A., Costantini, E., Forlani, S., Marsoni, M., Briani, F., Vannini, C., & Pesaresi, P. (2023). Perturbation of protein homeostasis brings plastids at the crossroad between repair and dismantling. PLOS Genetics, 19(7), e1010344. 10.1371/journal.pgen.1010344

Tadini, L., Peracchio, C., Trotta, A., Colombo, M., Mancini, I., Jeran, N., Costa, A., Faoro, F., Marsoni, M., Vannini, C., Aro, E. M., & Pesaresi, P. (2020). GUN1 influences the accumulation of NEP-dependent transcripts and chloroplast protein import in Arabidopsis cotyledons upon perturbation of chloroplast protein homeostasis. The Plant Journal, 101(5), 1198–1220. 10.1111/tpj.14585

Theron, C. W., Salcedo-Sora, J. E., Grixti, J. M., Møller-Hansen, I., Borodina, I., & Kell, D. B. (2024). Evidence for the Role of the Mitochondrial ABC Transporter MDL1 in the Uptake of Clozapine and Related Molecules into the Yeast Saccharomyces cerevisiae. Pharmaceuticals, 17(7), 938. 10.3390/ph17070938

Thomas, C., & Tampé, R. (2020). Structural and Mechanistic Principles of ABC Transporters. Annual Review of Biochemistry, 89(1), 605–636. 10.1146/annurev-biochem-011520-105201

van Wijk, K. J. (2015). Protein Maturation and Proteolysis in Plant Plastids, Mitochondria, and Peroxisomes. Annual Review of Plant Biology, 66(1), 75–111. 10.1146/annurev-arplant-043014-115547

van Wijk, K. J. (2024). Intra-chloroplast proteases: A holistic network view of chloroplast proteolysis. The Plant Cell, 36(9), 3116–3130. 10.1093/plcell/koae178

Verrier, P. J., Bird, D., Burla, B., Dassa, E., Forestier, C., Geisler, M., Klein, M., Kolukisaoglu, Ü., Lee, Y., Martinoia, E., Murphy, A., Rea, P. A., Samuels, L., Schulz, B., Spalding, E. P., Yazaki, K., & Theodoulou, F. L. (2008). Plant ABC proteins – a unified nomenclature and updated inventory. Trends in Plant Science, 13(4), 151–159. 10.1016/j.tplants.2008.02.001

Vishnepolsky, B., & Pirtskhalava, M. (2014). Prediction of Linear Cationic Antimicrobial Peptides Based on Characteristics Responsible for Their Interaction with the Membranes. Journal of Chemical Information and Modeling, 54(5), 1512–1523. 10.1021/ci4007003

Wagner, D., Przybyla, D., op den Camp, R., Kim, C., Landgraf, F., Lee, K. P., Würsch, M., Laloi, C., Nater, M., Hideg, E., & Apel, K. (2004). The Genetic Basis of Singlet Oxygen-Induced Stress Responses of *Arabidopsis thaliana*. Science, 306(5699), 1183–1185. 10.1126/science.1103178

Wolters, J. C., Abele, R., & Tampé, R. (2005). Selective and ATP-dependent translocation of peptides by the homodimeric ATP binding cassette transporter TAP-like (ABCB9). Journal of Biological Chemistry, 280(25), 23631–23636. 10.1074/jbc.M503231200

Yoo, S.-D., Cho, Y.-H., & Sheen, J. (2007). Arabidopsis mesophyll protoplasts: a versatile cell system for transient gene expression analysis. Nature Protocols, 2(7), 1565–1572. 10.1038/nprot.2007.199

Young, L., Leonhard, K., Tatsuta, T., Trowsdale, J., & Langer, T. (2001). Role of the ABC transporter Mdl1 in peptide export from mitochondria. Science, 291(5511), 2135–2138. 10.1126/science.1056957

Zhao, C., Haase, W., Tampé, R., & Abele, R. (2008). Peptide specificity and lipid activation of the lysosomal transport complex ABCB9 (TAPL). Journal of Biological Chemistry, 283(25), 17083–17091. 10.1074/jbc.M801794200

Zhao, H., Zhang, F., Wang, X., Liu, K., Zhang, L., Li, J., Kim, C., & Wang, L. (2025). The chloroplast translocon subunit TOC33 relays singlet oxygen-induced chloroplast-to-nucleus retrograde signaling in Arabidopsis. Molecular Plant, 18(10), 1706–1723. 10.1016/j.molp.2025.08.013

Zybailov, B., Coleman, M. K., Florens, L., & Washburn, M. P. (2005). Correlation of Relative Abundance Ratios Derived from Peptide Ion Chromatograms and Spectrum Counting for Quantitative Proteomic Analysis Using Stable Isotope Labeling. Analytical Chemistry, 77(19), 6218–6224. 10.1021/ac050846r

