## Supplementary figures and images for "Chloroplast ABC peptide transporters TAP1, NAP8, and ATH12 are essential for heat-induced peptide export and play a key role in thermotolerance in *Arabidopsis thaliana*"

### Supplemental figure 3

#### Fig S3 A

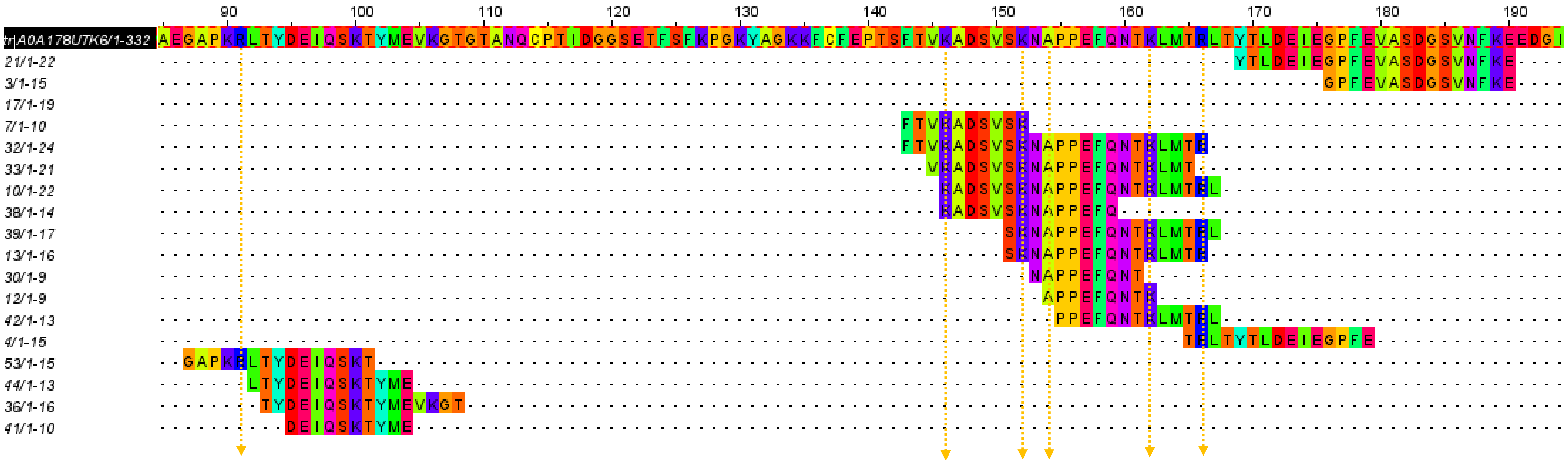

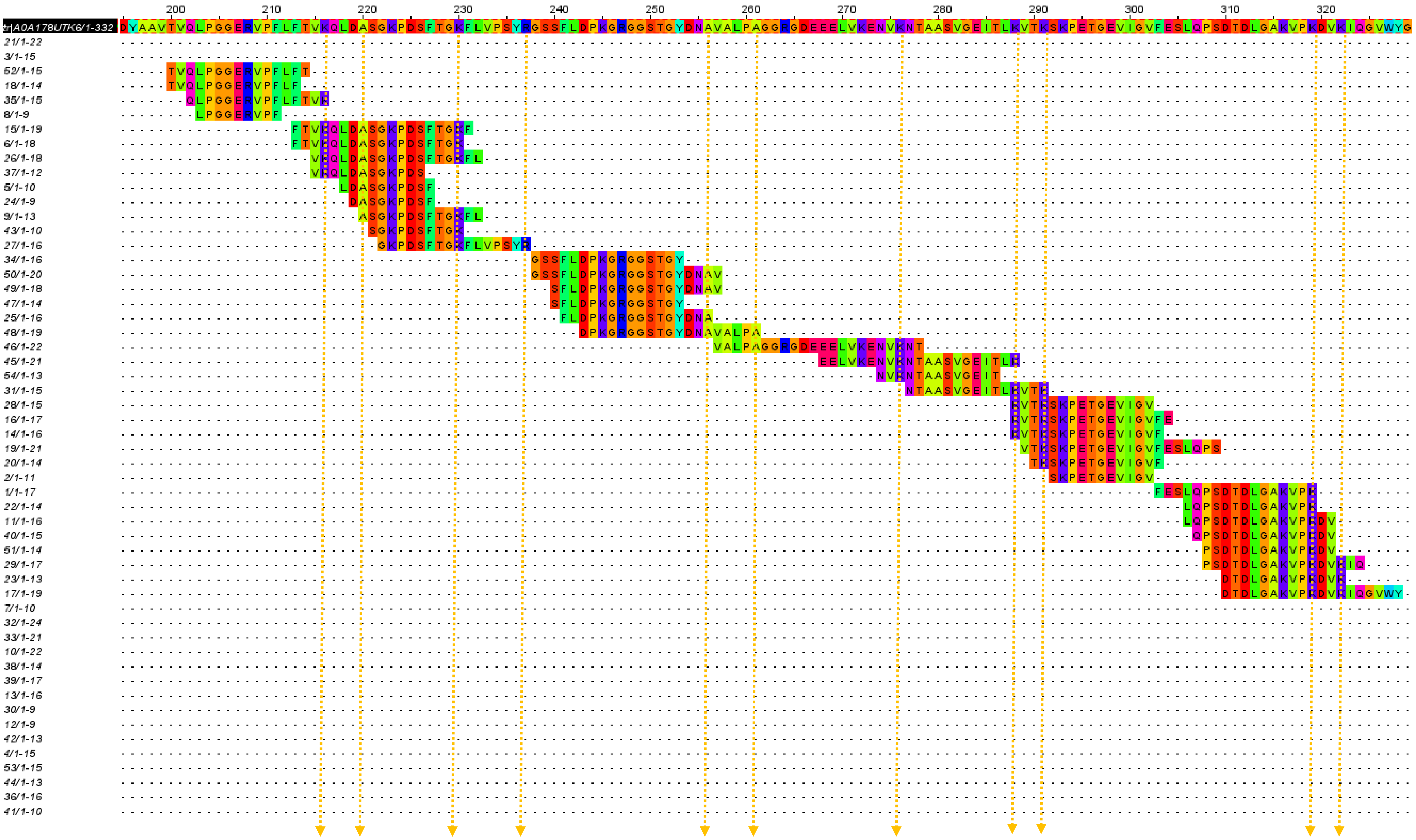

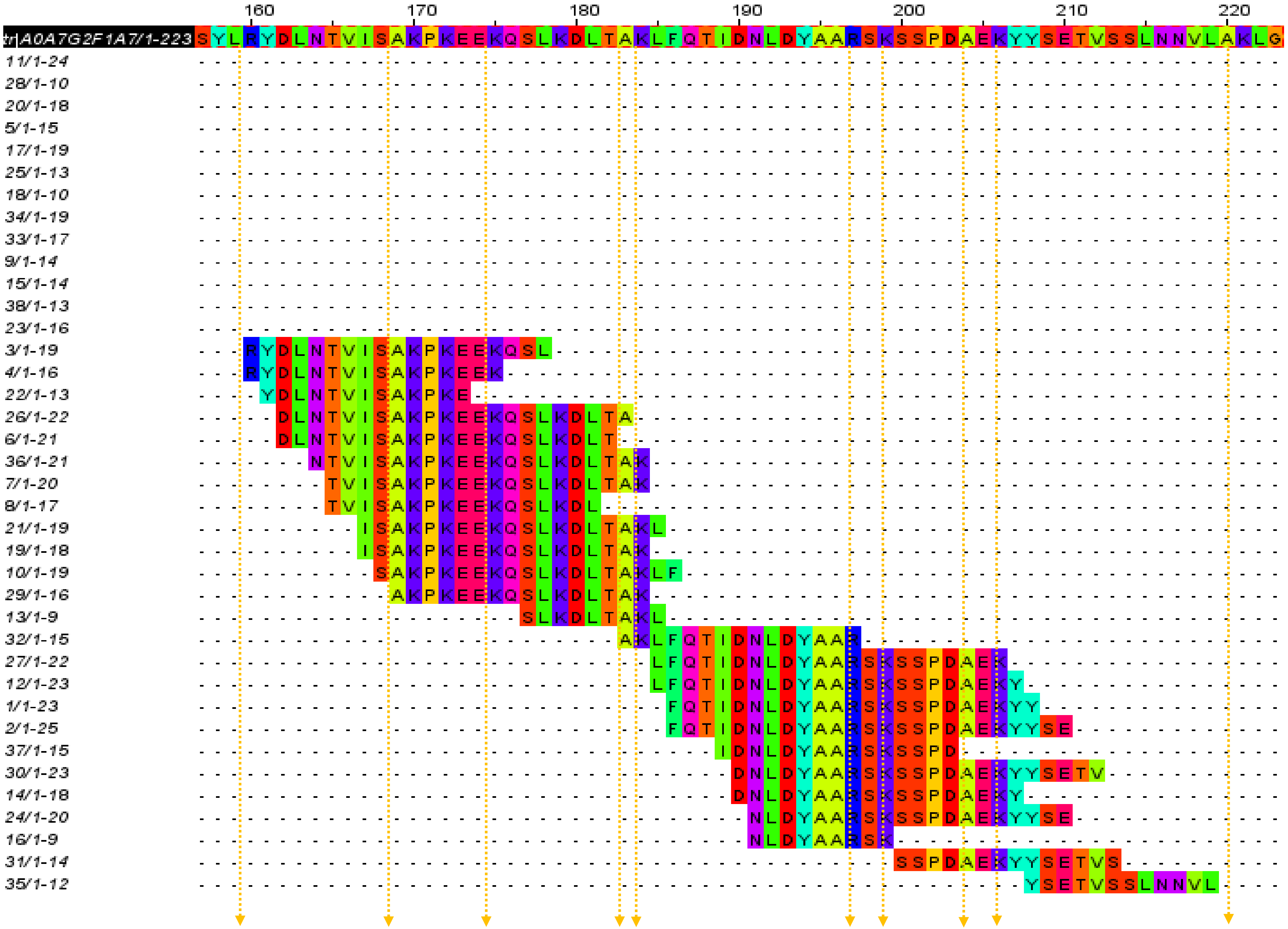

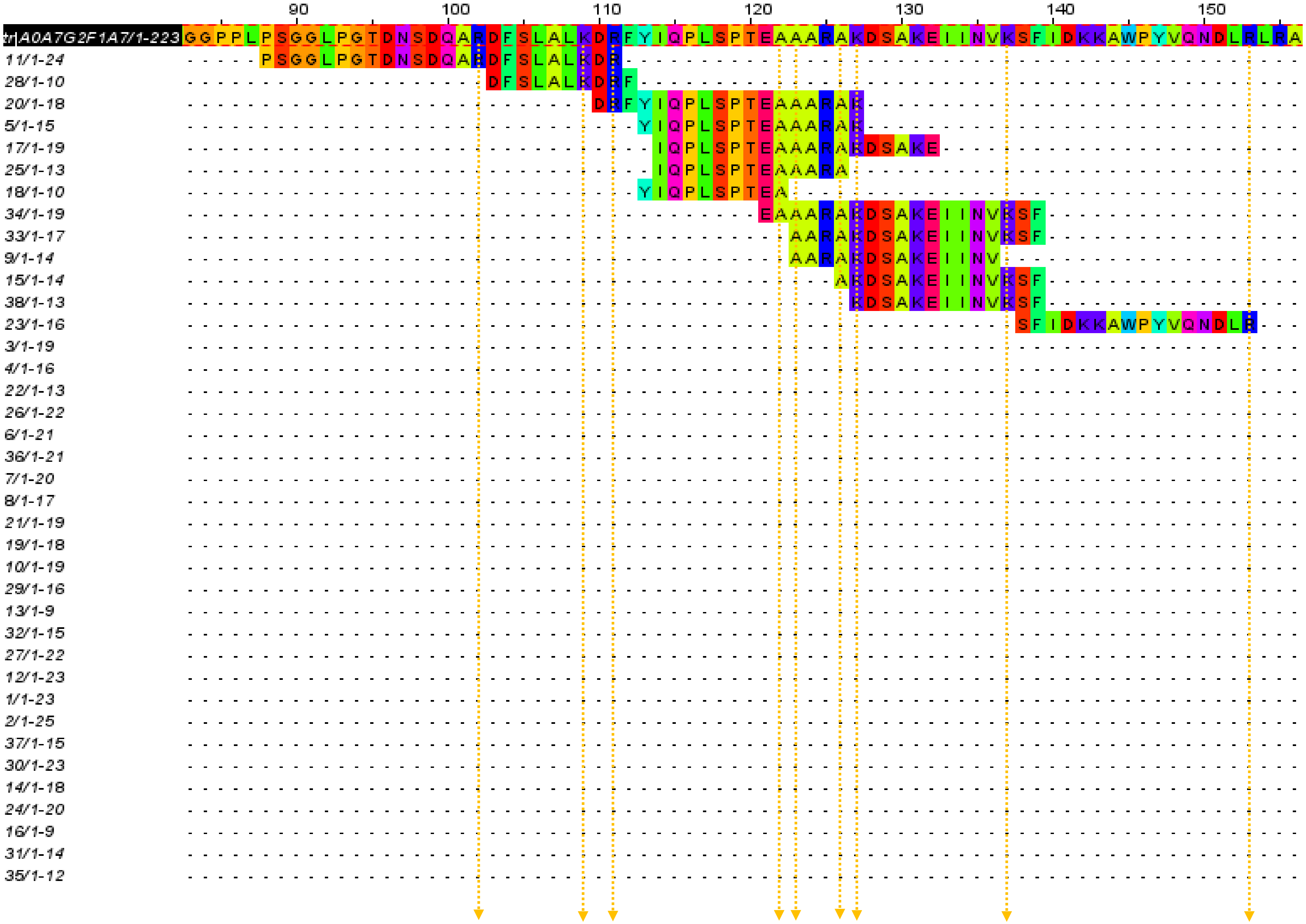

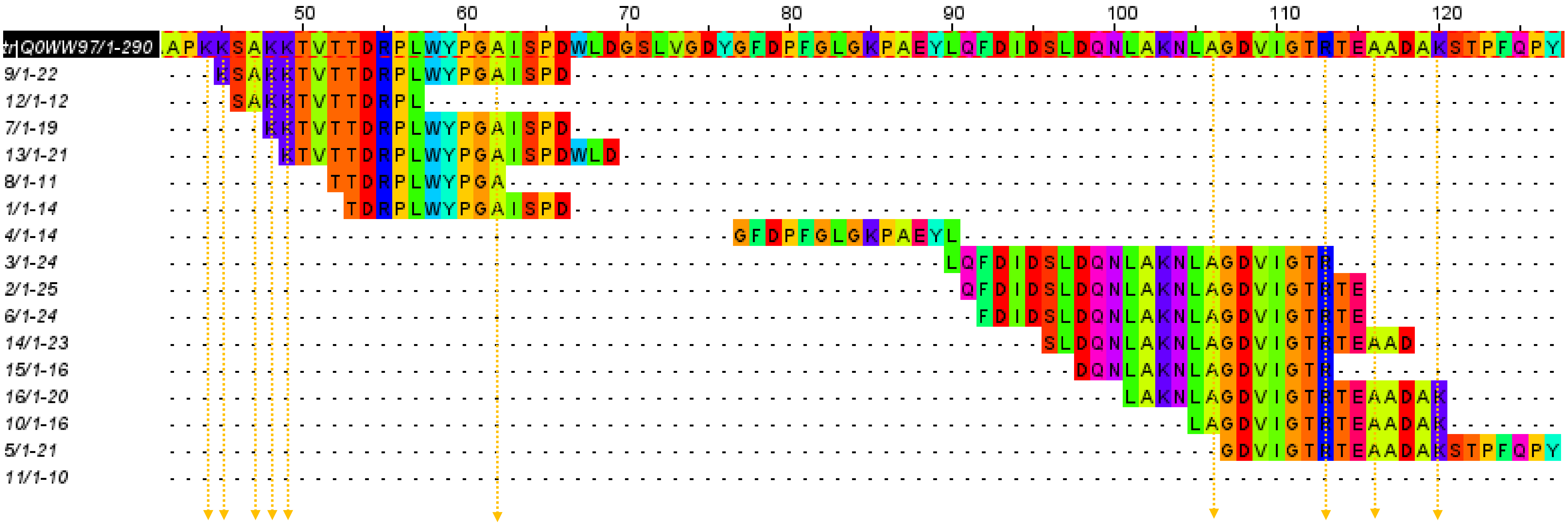

### Supplemental figures 1, 2, 4

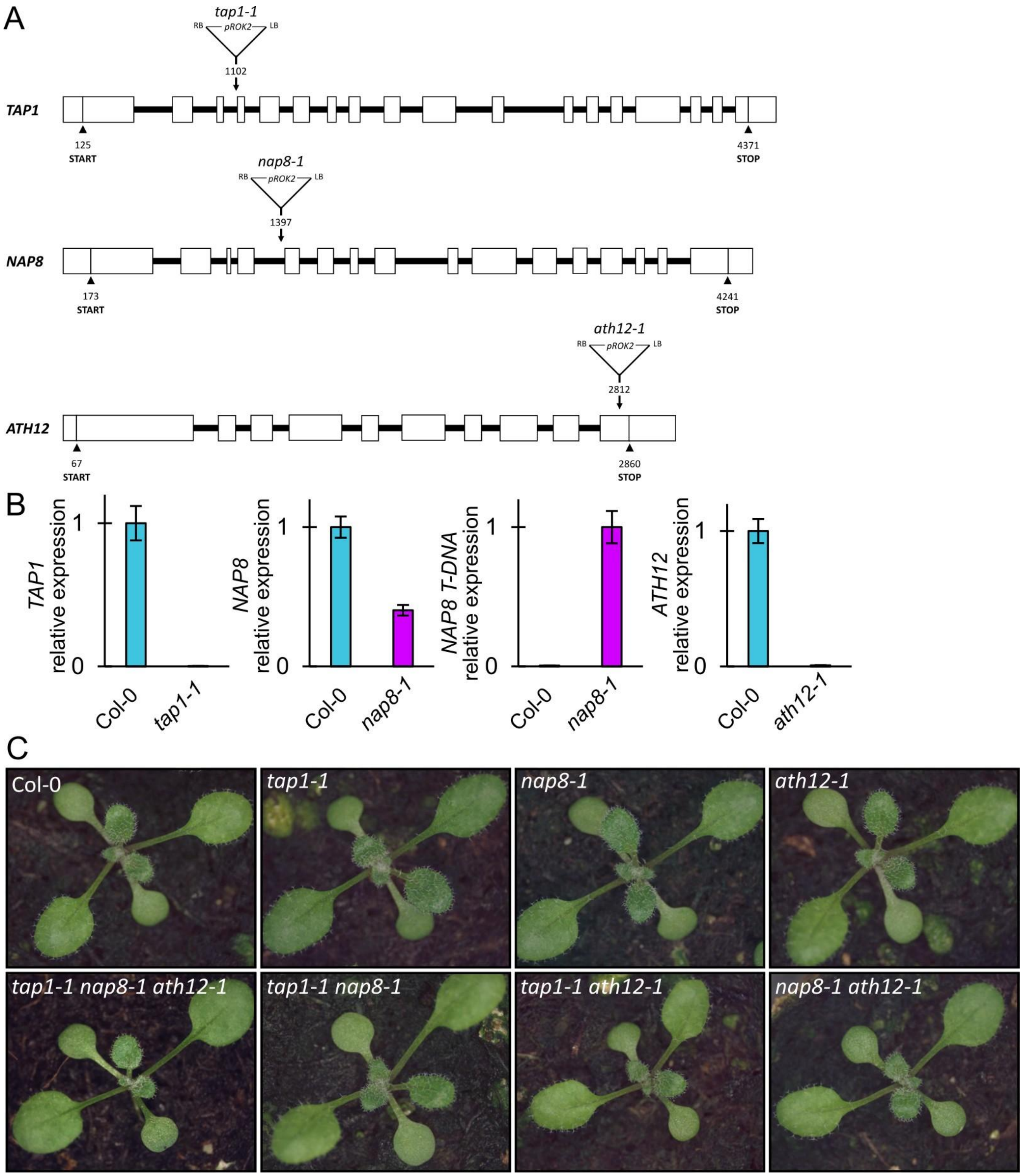

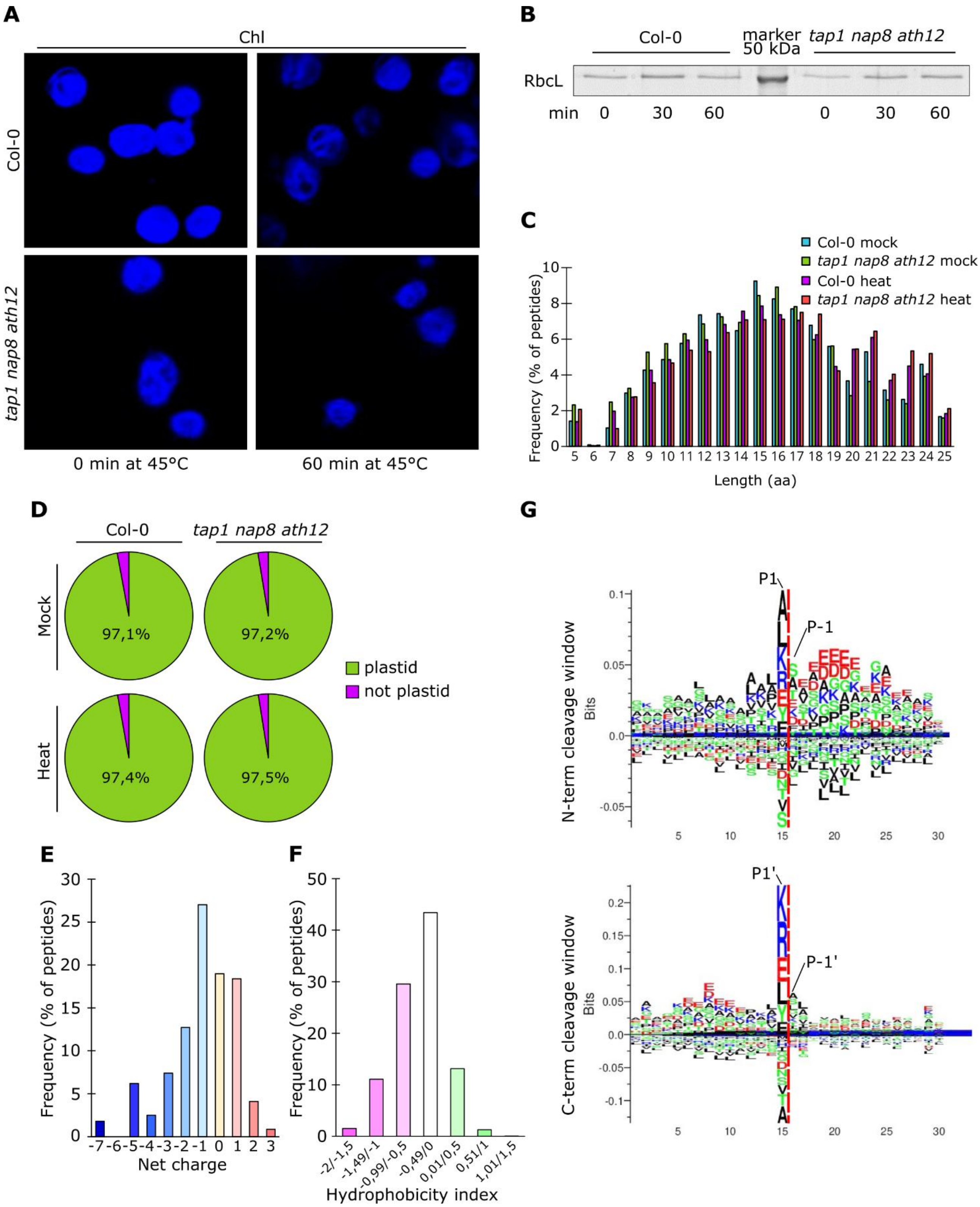

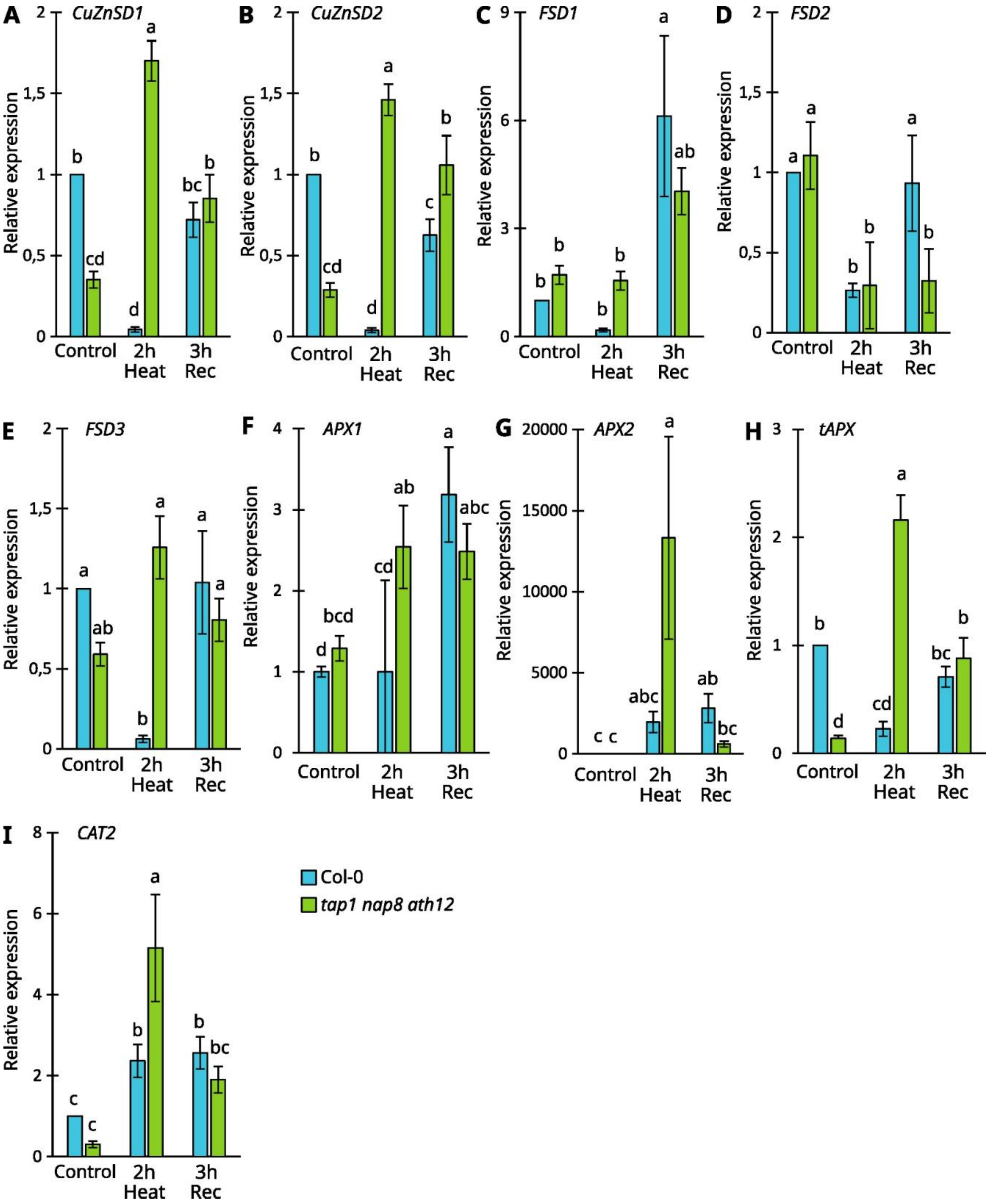
