## Supplemental table 7 for "Chloroplast ABC peptide transporters TAP1, NAP8, and ATH12 are essential for heat-induced peptide export and play a key role in thermotolerance in *Arabidopsis thaliana*"

| SOURCE PROTEIN | CORE SEQUENCE | NUMBER OF UNIQUE PEPTIDES WITH THE CORE SEQUENCE | SEQUENCE POSITIONS | LOCALIZATION |
|---|---|---|---|---|
| PSBO1 | PPEFQ | 9 | 155-159 | Lumenal |
| | GKPDS | 9 | 222-226 | Lumenal |
| | DTDLGAKVPK | 8 | 310-319 | Lumenal |
| | DPKGRGGSTGY | 6 | 243-253 | Lumenal |
| | DEIQSKT | 5 | 95-101 | Lumenal |
| | KADSV | 5 | 146-150 | Lumenal |
| | LPGGERVPF | 5 | 223-211 | Lumenal |
| | NTAASVGEIT | 3 | 277-286 | Lumenal |
| | EELVKENVKNT | 2 | 268-276 | Lumenal |
| PSBQ2 | AKPKE | 12 | 169-173 | Lumenal |
| | SLKDL | 10 | 177-181 | Lumenal |
| | NLDYAAR | 10 | 191-197 | Lumenal |
| | SSPD | 9 | 200-203 | Lumenal |
| | IQPLSPTEA | 6 | 114-122 | Lumenal |
| | AARAK | 6 | 122-127 | Lumenal |
| | KDSAKEIINV | 5 | 127-136 | Lumenal |
| | DFSLALAKDR | 3 | 103-111 | Lumenal |
| LHCB4 | GDVIGTR | 9 | 107-113 | Stromal |
| | TDRPL | 6 | 53-57 | Stromal |
